# How did evolution halve genome size during an oceanic island colonization?

**DOI:** 10.1101/2025.06.14.659697

**Authors:** Vadim A. Pisarenco, Adrià Boada-Figueras, Marta Olivé-Muñiz, Paula Escuer, Nuria Macías-Hernández, Miquel A. Arnedo, Pablo Librado, Alejandro Sánchez-Gracia, Sara Guirao-Rico, Julio Rozas

**Author notes:** Equal contribution (Vadim A. Pisarenco, Adrià Boada-Figueras, Marta Olivé-Muñiz). corresponding authors Julio Rozas, Sara Guirao-Rico.

## Abstract

Red devil spiders of the genus *Dysdera* colonised the Canary Islands and underwent an extraordinary diversification. Notably, their genomes are nearly half the size of their mainland counterparts (∼1.7 *vs.* ∼3.3 Gb). This offers a unique model to solve long-standing debates regarding the roles of adaptive and non-adaptive forces on shaping genome size genome size evolution. To address these, we conducted comprehensive genomic analyses based on three high-quality chromosome-level assemblies, including two newly generated ones. We find that insular species experienced a reduction in genome size, affecting all genomic elements, including intronic and intergenic regions, with transposable element (TE) loss accounting for most of this contraction. Additionally, autosomes experienced a disproportionate reduction compared to the X chromosome. Paradoxically, island species exhibit higher levels of nucleotide diversity and recombination, lower TE activity in recent times, and evidence of intensified natural selection, collectively pointing to larger long-term effective population sizes in species from the Canary Islands. Overall, our findings align with the non-adaptive mutational hazard hypothesis, supporting purifying selection against slightly deleterious DNA and TE insertions as the primary mechanism driving genome size reduction.

## Introduction

Understanding the evolutionary processes underlying genome size variation is a central and long-standing question in evolutionary biology (Linquist et al. 2015; Wright 2017; Blommaert 2020; Galtier 2024). genome size exhibits remarkable diversity across eukaryotes and occasionally within and among closely related species (Huang et al. 2014; Naville et al. 2019; Stelzer et al. 2023). Two main mechanisms contribute to rapid increases in genome size: changes in ploidy, and transposable element (TE) mobilisation (Sproul et al. 2023). While whole genome duplication (WGD) is widespread in plants, it only occurs sporadically in animals (Li et al. 2021; Galtier 2024). Moreover, certain lineages, such as birds (Kapusta et al. 2017), exhibit a relatively stable genome size over time. Despite their potential deleterious effects, WGDs occasionally confer selective advantages as a source of new genetic material and functional innovation (Ohno 1970). This also applies to TE insertions, which can occasionally increase individual fitness in front of sudden environmental changes (Casacuberta and González 2013; Baduel et al. 2018; Dubin et al. 2018; Van de Peer et al. 2021).

Differences in repetitive DNA content, particularly of TEs, accounts for the vast majority of genome size variation across eukaryotes (Wright 2017; Arkhipova 2018). However, the evolutionary forces and mechanisms behind genome size variation remain hotly debated (Whitney et al. 2011; Lynch et al. 2023; Galtier 2024). Several hypotheses have been proposed to explain genome size variation, many of which are not mutually exclusive (Blommaert 2020). Non-adaptive hypotheses posit that genome size is predominantly shaped by negative selection and neutral processes such as mutation and genetic drift. These include the mutational equilibrium (Petrov 2001; Petrov 2002) and the mutational hazard (Lynch and Conery 2003; Lynch 2011; Smith 2016) hypotheses. The mutational equilibrium hypothesis provides the simplest model of genome size evolution, proposing that expansions occur through gene duplications and transposition bursts, while contractions result from small deletions scattered along the chromosomes. A variation of this hypothesis, the so-called accordion model, suggests instead that large-scale segmental deletions account for most DNA loss (Kapusta et al. 2017). In contrast, the mutational hazard hypothesis suggests that the fate of DNA gains, such as gene duplications and TEs expansions, is governed by the mutation rate and the effective population size (*N*_e_). While the per-site mutation rate is expected to be lower in longer genomes (Lynch and Conery 2003; Lynch 2011), a larger *N*_e_ increases the efficacy of selection, more effectively purging slightly deleterious DNA gains and losses.

In contrast, adaptive hypotheses position genome size as a direct target of natural selection, suggesting an associated fitness value, for instance, through cell size changes (Gregory and Hebert 1999). Cell size influences adaptive life history traits, such as body size, division rate or developmental time (Gregory 2004; Cavalier-Smith 2005; Tsukaya 2013; Liedtke et al. 2018). Other influential adaptive hypotheses include the genome streamlining hypothesis (Hessen et al. 2010; Stelzer et al. 2023), which suggests that increased selective pressures tend to shrink genome size in phosphorus or nitrogen-deficient environments, two essential components of DNA. Another hypothesis, proposed by (Yin et al. 2018), suggests a direct link between genome size and the evolution of self-fertility.

Unravelling the evolutionary forces shaping genome size variation requires an appropriate study model to evaluate whether the observed patterns and levels of DNA gain and loss are best explained by adaptive hypotheses, align with the mutational patterns predicted by the neutral theory (supporting the mutational equilibrium hypothesis and similar models), or correlate with the *N*_e_, which can be considered as a proxy for the efficacy of selection, as predicted by the mutational hazard hypothesis. Here, we used the island radiation of the spider genus *Dysdera* in the Canary Islands as a natural laboratory to study genome size evolution. This genus underwent an important evolutionary diversification in the Canary Islands, with almost 50 endemic species (14% of the total described) emerging since the formation of this Macaronesian archipelago (Arnedo et al. 2001; Arnedo et al. 2007; Macias-Hernandez et al. 2016; Řezáč et al. 2021; Bellvert et al. 2023; Bellvert, Pollock, et al. 2024). Current phylogenetic evidence suggests that the Canary Islands endemics of the genus *Dysdera* form a clade that is sister to most species currently inhabiting the western Mediterranean basin, including northern Africa and the oceanic archipelagoes of Madeira and the Azores. These latter groups are, in turn, related to species from the Middle East and Central Asia, the *asiatica* and *lata* groups *sensu* (Deeleman-Reinhold and Deeleman 1988). The most basal split within the genus would involve the former clade and a group of species inhabiting the Eastern Mediterranean (the *logirostris* and *ninni* groups) (Adrián-Serrano et al. 2021; Crespo, Isamberto, et al. 2021; Bellvert, Dimitrov, et al. 2024). An unexplained feature of the island species is that their genome size is approximately half that of their mainland counterparts (Escuer et al. 2022), while maintaining similar lifestyles (Řezáč et al. 2018; Bellvert et al. 2023). To investigate this, we generate two new genome chromosome-level assemblies, one for an insular (*D. tilosensis* Wunderlich, 1992; haploid genome size ∼1.7 Gb) and another for a mainland (*D. catalonica* Řezáč, 2018; haploid genome size 3.3 Gb) species, and analyse them alongside a third assembly, from the island species *D. silvatica* (haploid genome size ∼1.7 Gb (Escuer et al. 2022)), and incorporating in our study newly generated DNA resequencing data.

## Results

### Genome size and chromosome-level assemblies

Using flow cytometry, we determined that the genomes of the six surveyed Canary Island species approximately span 1.7 Gb, whereas those from mainland and Madeira species are around 3 Gb (Fig. 1a). We generated two new chromosome-level assemblies in this study, one for a mainland and another for an insular *Dysdera* species, namely *D. catalonica* and *D. tilosensis*, respectively (Figs. 1b and c; supplementary tables S1 and S2). Both genome assemblies were consistent with our flow cytometry estimates (3.3 Gb and 1.7 Gb, respectively; Fig. 1a), exhibited high completeness (93.1% and 94.9% complete BUSCO genes using the Arthropoda dataset, respectively; supplementary tables S3) and continuity (N50 values of 604.5 Mb and 210.8 Mb, respectively; Table 1). The number of large scaffolds matched the haploid chromosome number previously described in Řezáč et al. (2018) and in Escuer et al. (2022) (Fig. 1d). In total, 89.9% and 96.7% of the *D. catalonica* and *D. tilosensis* assembled sequences are contained within their chromosome-level scaffolds (Table 1). Paradoxically, the species with the largest genome only includes four autosomes and the X chromosome (Řezáč et al. 2018), whereas the halved *D. tilosensis* genome has six autosomes and the X chromosome.

**Fig. 1.**
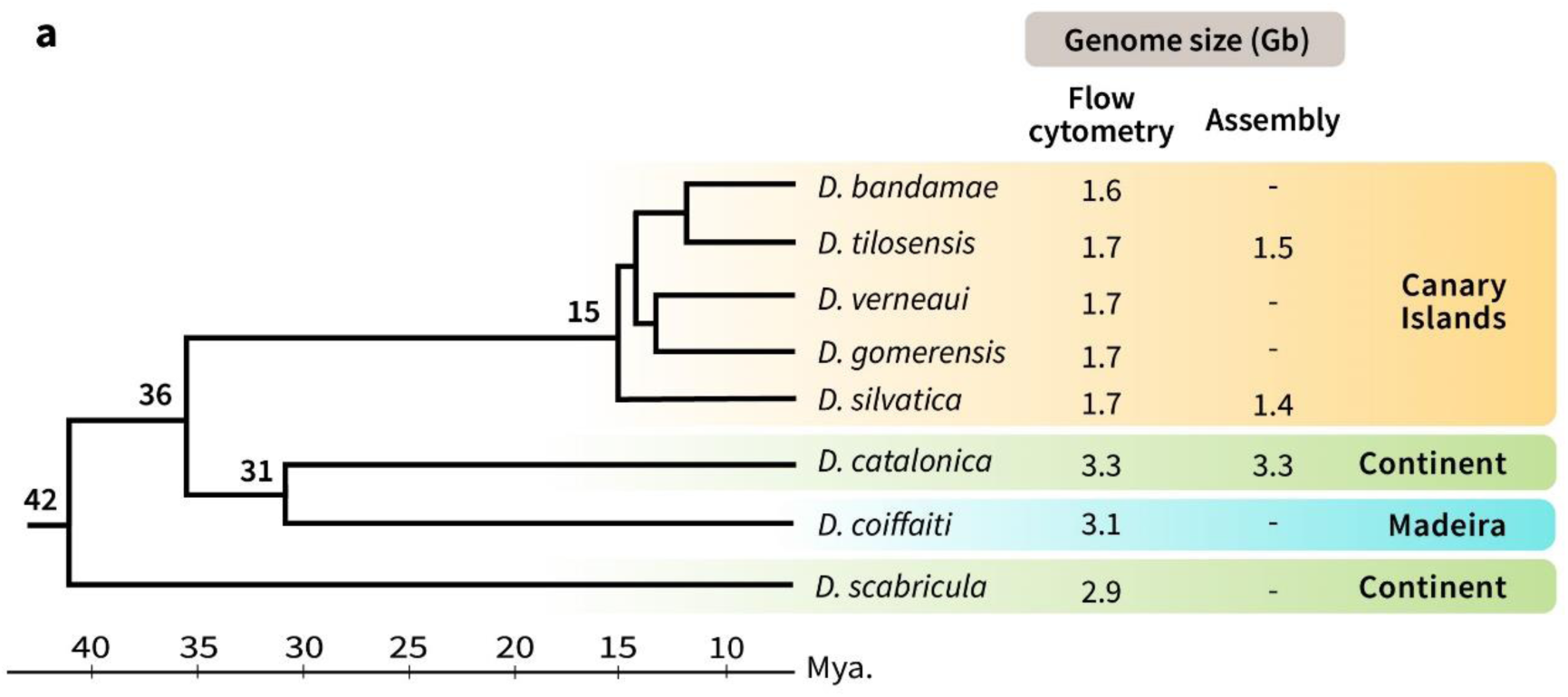
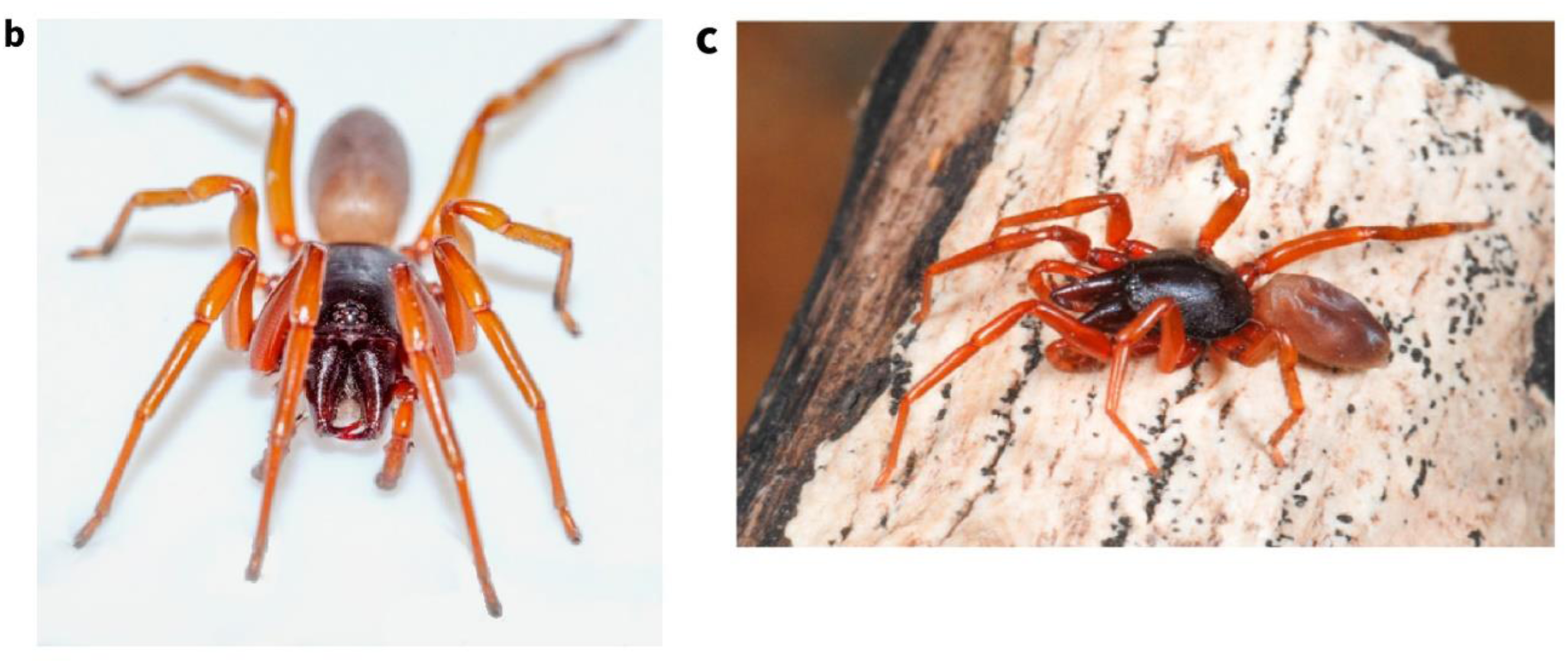
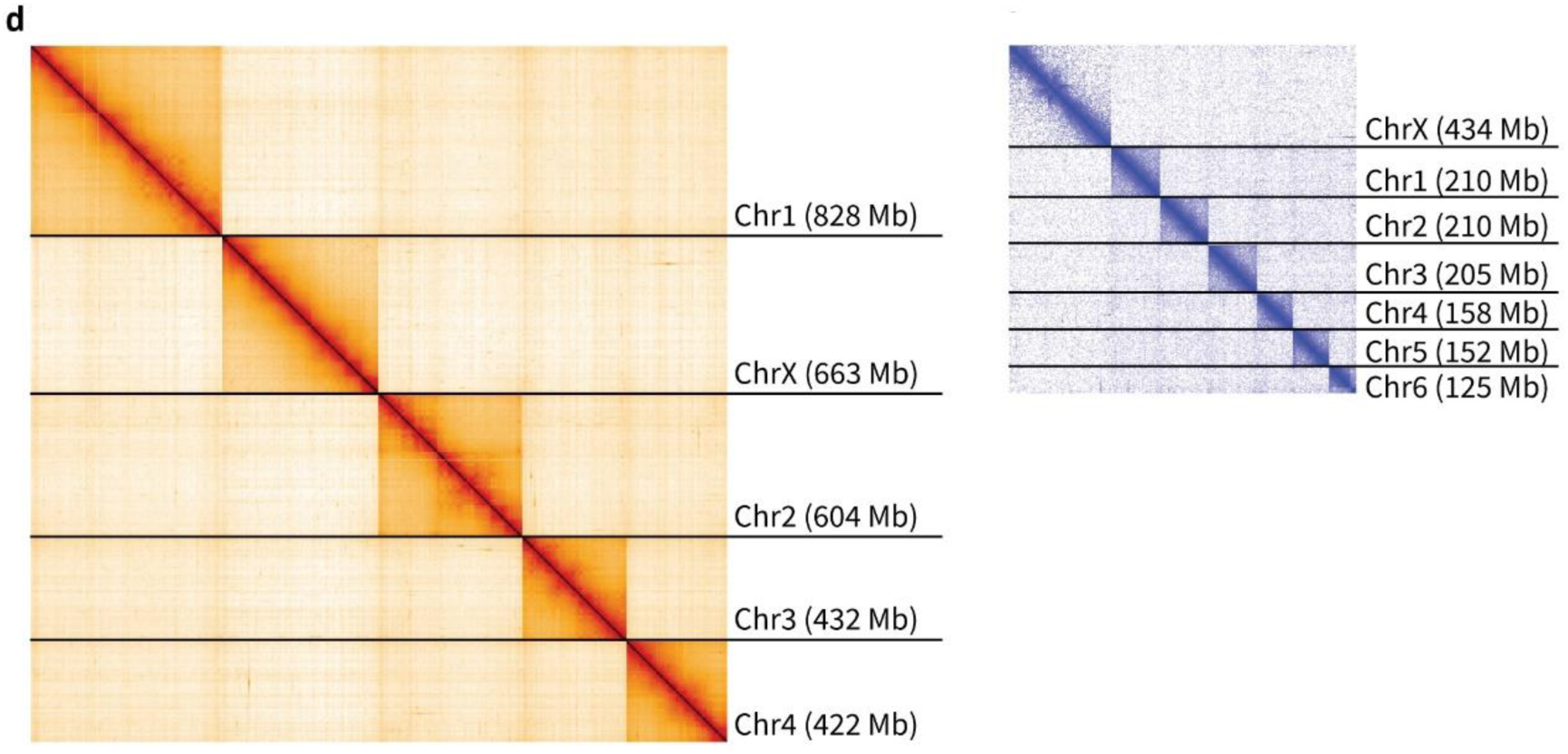
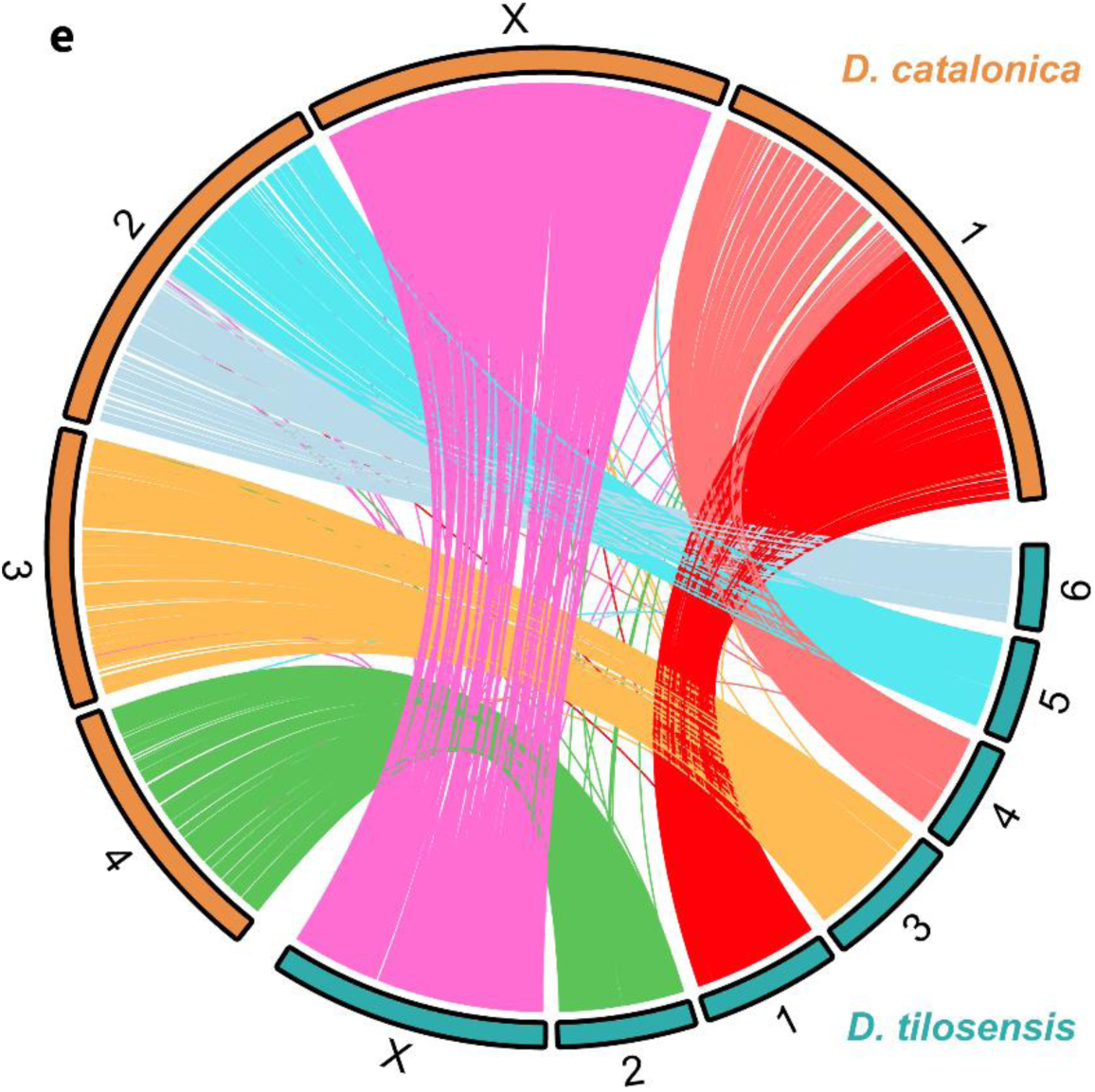
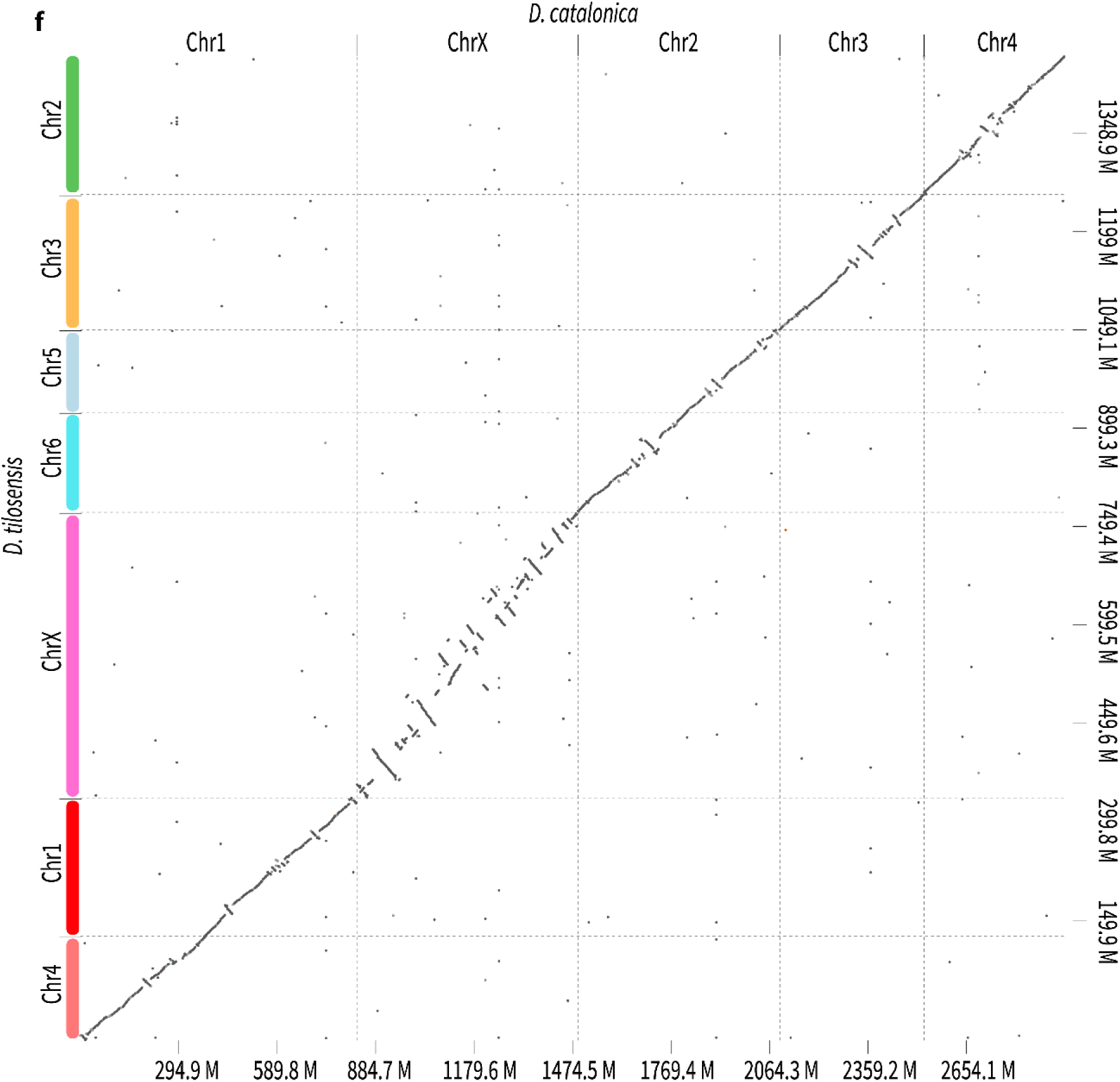
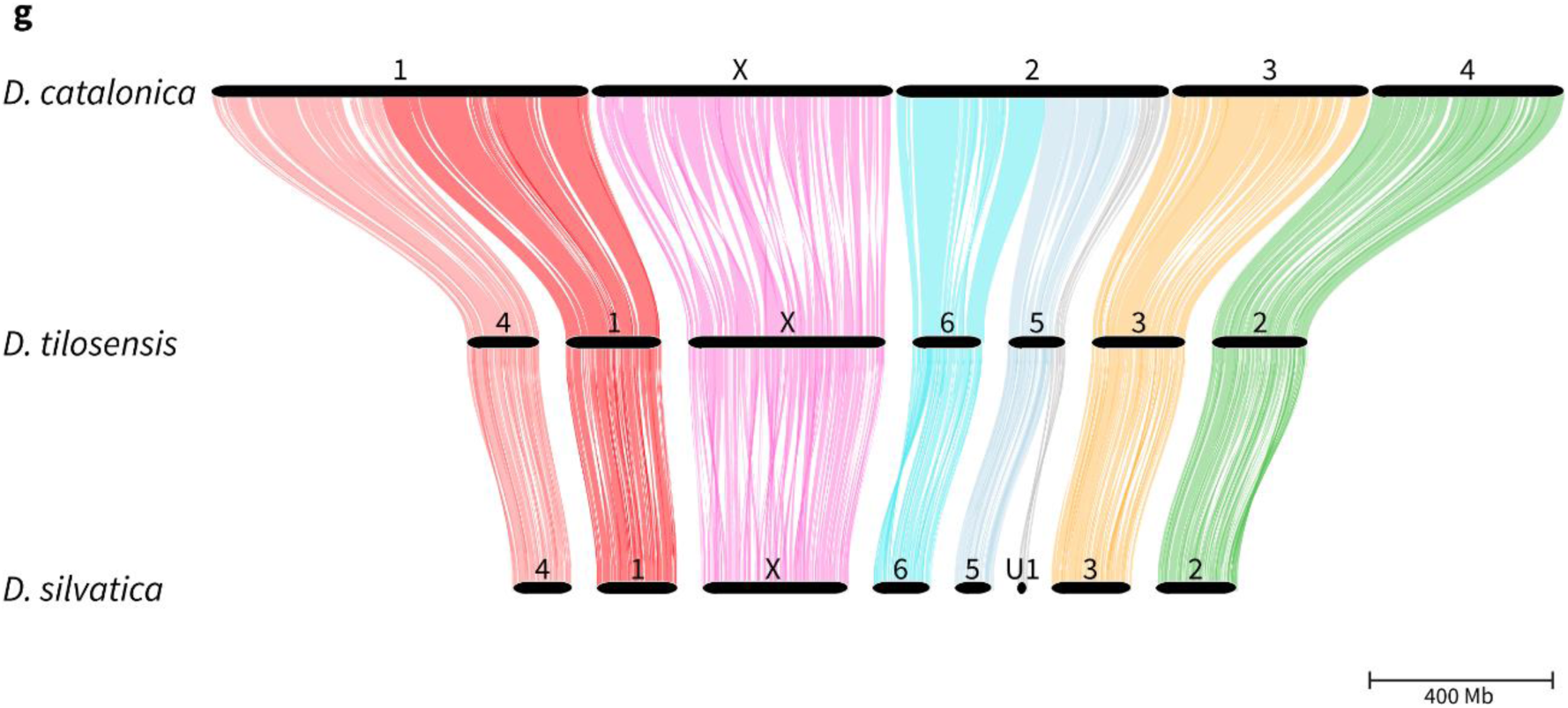
a) Phylogenetic relationship and divergence times of insular and continental species, representative of the *Dysdera* genus, based on Crespo et al. (2021). These include representatives from all major clades within the genus. Numbers at the nodes indicate divergence ages (in Mya). b) Specimens of *D. catalonica* and c) *D. tilosensis* (Photo credit: Marc Domènech and Pedro Oromí, respectively). d) Genome-wide OmniC contact maps of the chromosome-level assemblies of *D. catalonica* (left) and *D. tilosensis* (right). Contact maps are scaled by assembly size. e) Chromosomal synteny (CIRCOS plot) obtained by mapping the genome of *D. tilosensis* (query) to *D. catalonica* (target) using minimap2. The outer label delineates the autosome number and the X chromosome. f) D-GENIES plot based on the minimap2 alignments of *D. catalonica* and *D. tilosensis* genomes, showing a highly conserved synteny with no signs of WGD. g) The GENESPACE syntenic map (‘riparian plot’) of orthologous regions among the three *Dysdera* species.

**Table 1.** Genome assembly and annotation statistics.

|  | *D. catalonica* | *D. tilosensis* |
| --- | --- | --- |
| Assembly size (bp) | 3,279,770,110 | 1,549,768,629 |
| GC Content (%) | 35.50% | 34.78% |
| Number of scaffolds | 4,461 | 3,756 |
| Longest scaffold (Mb) | 827.9 | 434.8 |
| Scaffold N50 (Mb) / L50 | 604.5 / 3 | 210.8 / 3 |
| Chromosomes (a) & Sequence in chromosomes (%) | 5 / 89.91% | 7 / 96.71% |
| Genome annotation |  |  |
| Protein-coding genes | 47,753 | 22,699 |
| Genes with SBS evidence | 23,223 | 16,922 |
| Functionally annotated | 27,913 | 15,758 |
| Repetitive regions (%) | 73.6% | 64.9% |
| Orthogroups |  |  |
| Orthogroups (b) | 16,893 | 16,934 |
| Genes in orthogroups (c) | 43,046 | 20,908 |
| Orthogroups 1:1 (d) | 7,547 | 7,547 |

(a) Haploid number of chromosomes.
(b) Number of orthogroups identified by Broccoli and Orthofinder.
(c) Number of genes in orthogroups identified by Broccoli.
(d) Number of orthogroups excluding chimeric cases and overlapping genes.

Structural annotation predicts that the genome of *D. catalonica* encodes 47,753 protein-coding genes, more than twice the 22,699 identified in *D. tilosensis*. This marked difference likely stems from two main factors. First, it is well known that annotation tools often generate fragmented gene models in species with large intronic distances, particularly in non-model organisms such as *D. catalonica,* which can artificially inflate gene counts. To address this issue, we conducted two complementary analyses: (1) estimation of the number of high-confidence gene sets (SBS gene sets), and (2) quantification of the total number of identified exons, a metric less sensitive to annotation artifacts than overall gene counts. While both analyses reduced the discrepancy, *D. catalonica* still shows over 35% more high-confidence genes and exons than *D. tilosensis* (Table 1; Supplementary Table 4). A second key insight emerges from the analysis of orthogroups: despite the substantial difference in gene counts, the number of orthogroups is nearly identical between the two species (16,893 in *D. catalonica* and 16,934 in *D. tilosensis*; Table 1; Supplementary Table 4). This suggests that the remaining disparity is primarily due to a higher number of paralogous gene copies per orthogroup in the mainland lineage.

As previously described for another Canarian endemic species, *D. silvatica* Schmidt, 1981 (Escuer et al. 2022), a significant portion of the new surveyed genomes is composed of repetitive elements (REs), mostly TEs, which span 72.5% and 63.5% of the *D. catalonica* and *D. tilosensis* autosomes, and 76.1% and 68.4% of their X chromosomes (Fig. 2a, Table 2; supplementary tables S5 and S6). Globally, DNA transposons (class II TEs) occupy higher genome proportions than retrotransposons (class I TEs), representing 42.0% and 28.2% of the *D. catalonica* genome, and 39.3% and 22.6% of that of the *D. tilosensis* (supplementary table S5). While retrotransposon composition, such as LTRs, LINEs, and SINEs, varies markedly between species (see below) (Fig. 2b), the overall TE proportion remains quite similar in mainland and Canarian species, despite their remarkable genome size differences.

**Fig. 2.**
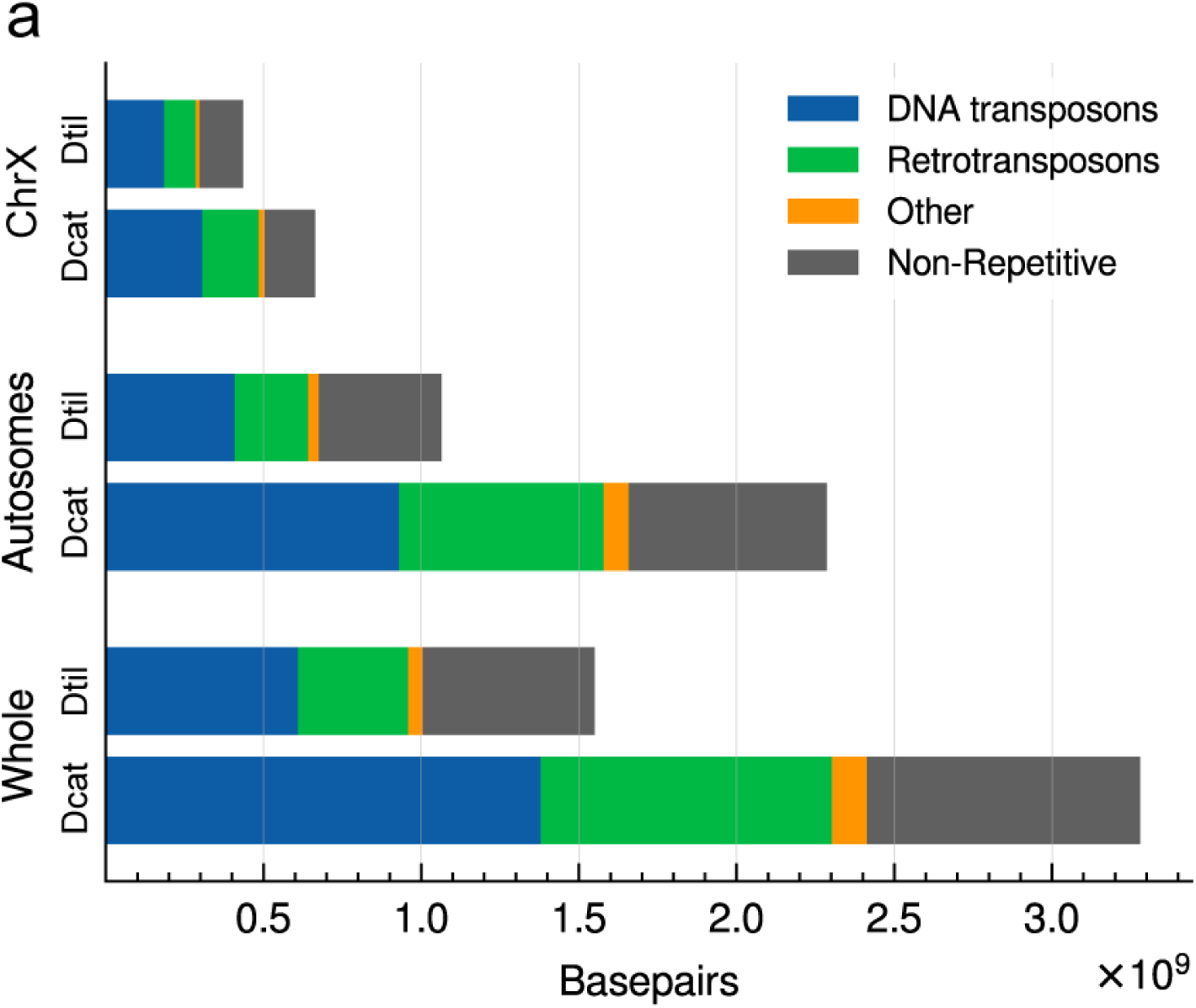
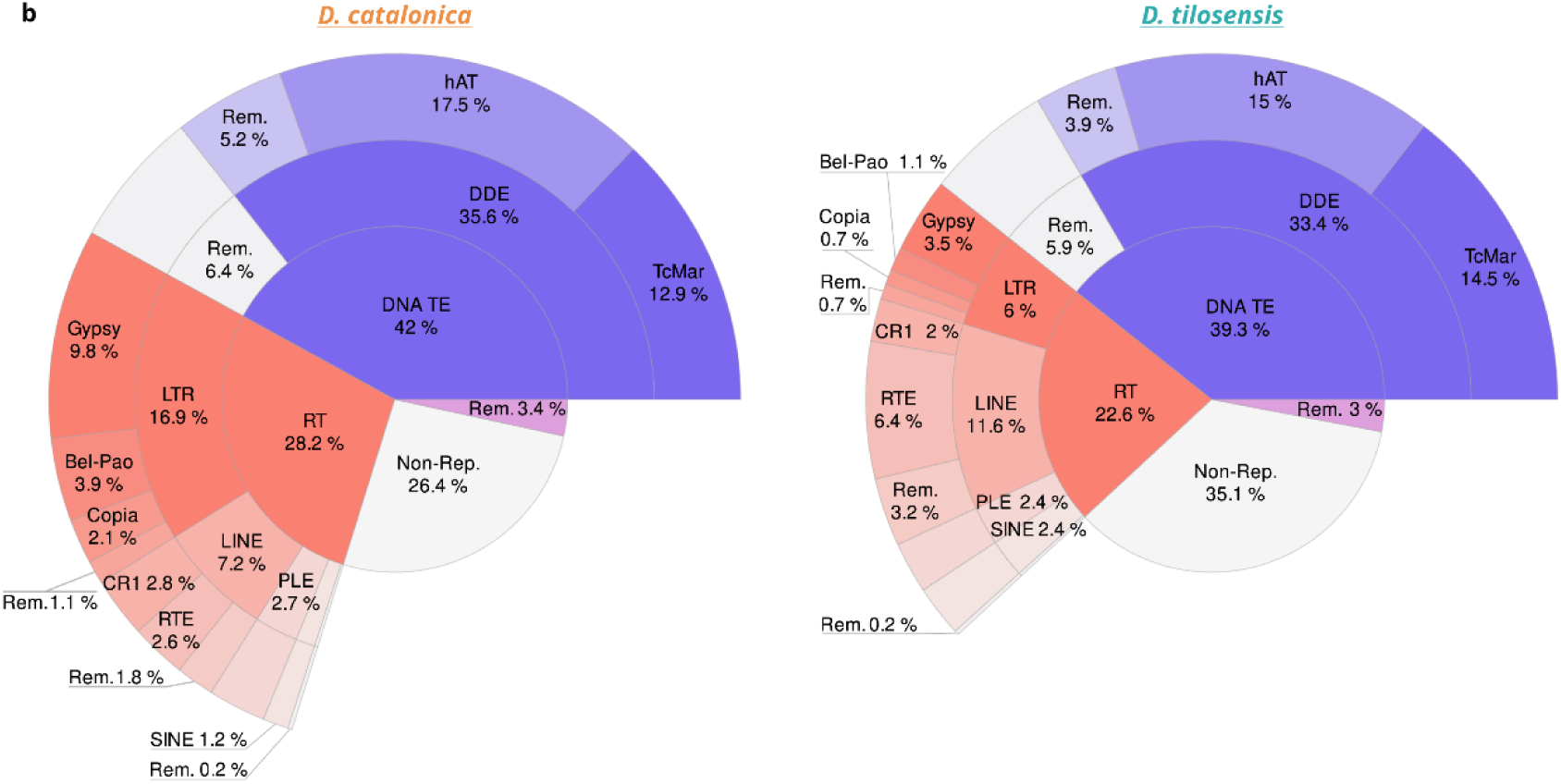
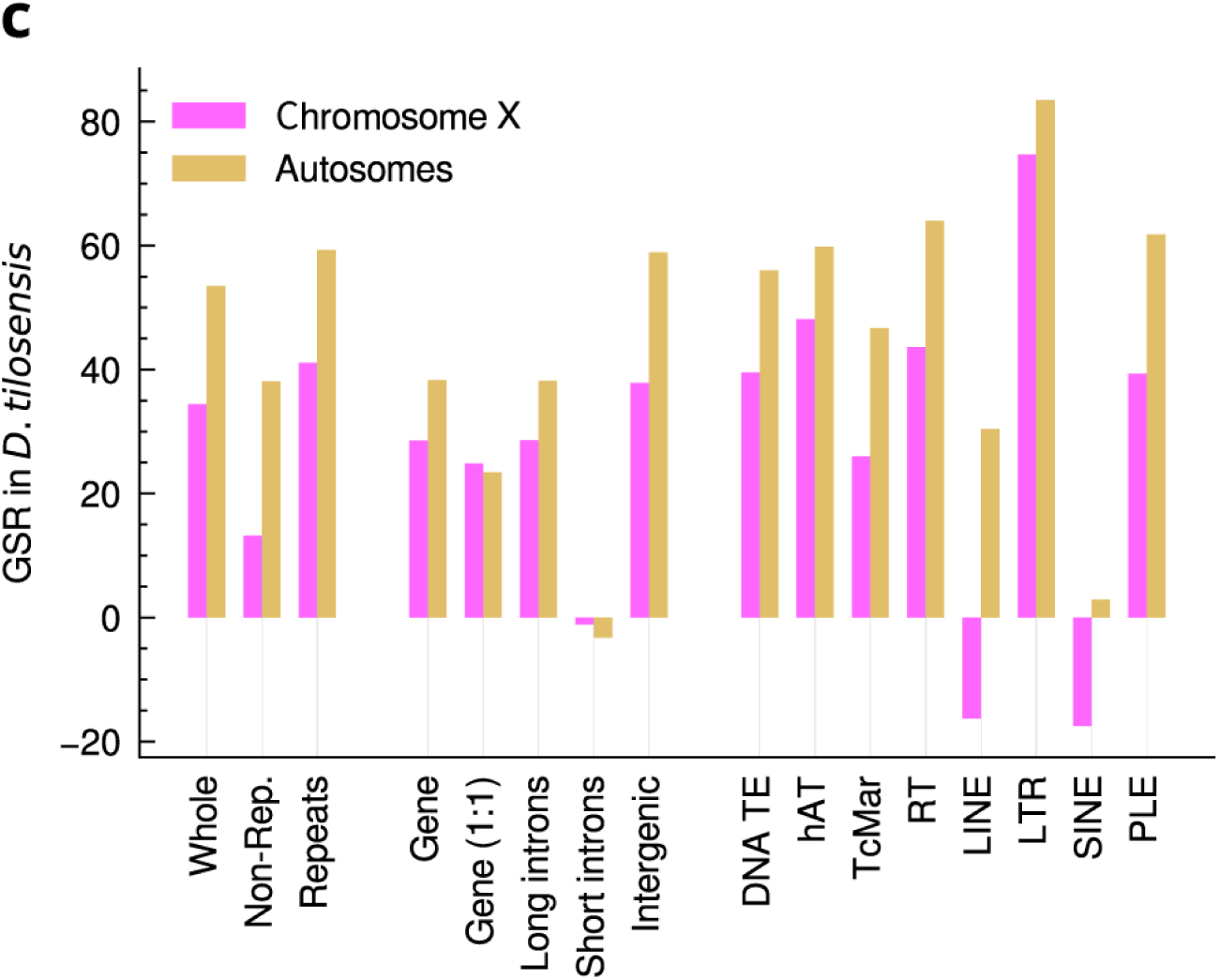
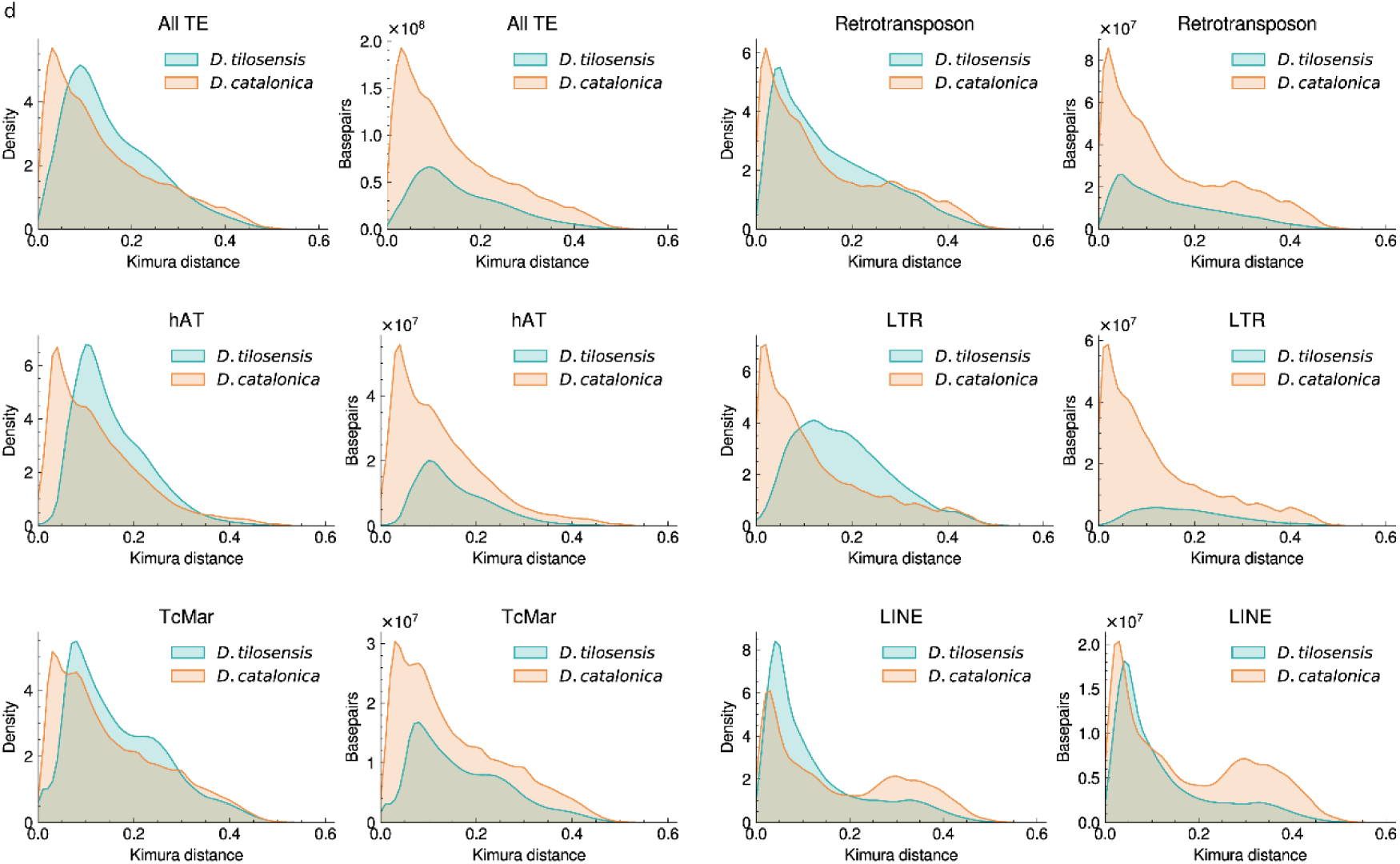
a) Size (in bp) of the repetitive and non-repetitive fraction of the genome assembly. The ‘whole genome’ bars include the complete genome information (large and small scaffolds). b) TE distribution in *D. catalonica* and *D. tilosensis. Rem.* stands for the remaining repeat categories. c) Percentage of genome size reduction (GSR) in *D. tilosensis* across several genomic features, comparing the X chromosome and autosomes (excluding minor scaffolds). “Whole”, “Non-Rep.” and “Repeats” stand for the whole genomic subset, the non-repetitive and repetitive fraction, respectively (see also supplementary table 5). “Gene” and “Gene (1:1)” indicate the total number of genes and one-to-one orthologs, including introns. “Long introns” refer to intronic regions longer than 300bp, whereas “Short introns” include regions of 300 bp or less. “Intergenic”, intergenic regions detailed in supplementary table 9. “DNA TE”, “hAT”, “TcMar”, “RT”, “LINE”, “LTR”, “SINE”, “PLE”, stand for DNA transposons, hAT superfamily, TcMar superfamily, retrotransposons, LINE order, LTR order, SINE order and Penelope order, respectively. d) Kernel density estimates for Kimura divergence of different TE groups. The Kimura genetic distances are adjusted for the CpG sites.

**Table 2.**
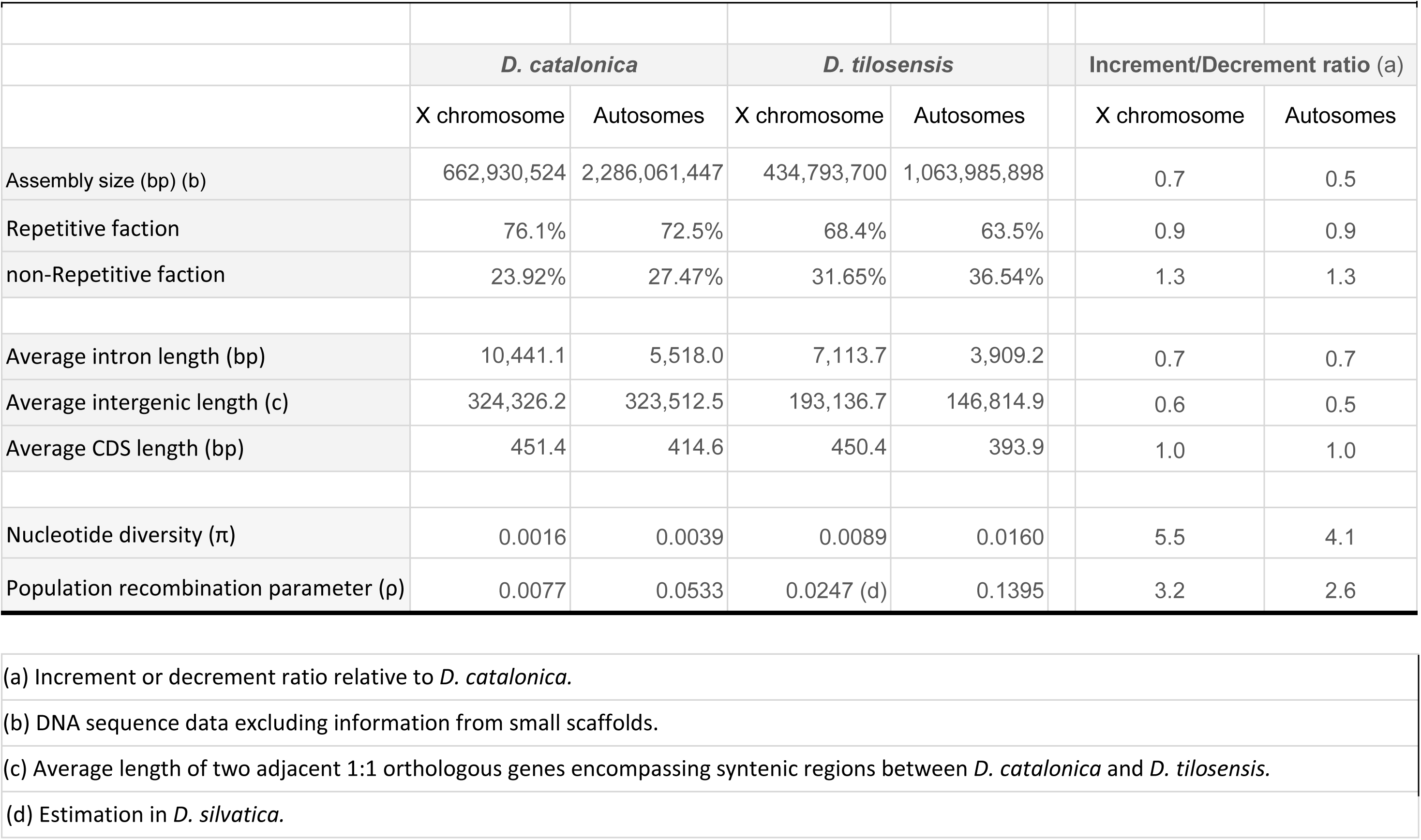
Genome features across autosomes and X chromosomes.

### Evidence for an ancestral genome size reduction in the lineage leading to *Dysdera* species from the Canary Islands

Multiple lines of evidence support that the nearly twofold difference in genome size between mainland and insular species resulted from a genome contraction process in the insular species, rather than a WGD in the continental lineage (Fig. 1e-g; supplementary fig. S1). Firstly, genome synteny analyses reveal high genome collinearity across species, and no trace of large duplicated segments with high intragenomic collinearity, typical of recent WGD events (Van de Peer 2004). Indeed, we only observed two major chromosomal rearrangements, indicative of two fusion/fission events (Fig. 1e-g). Secondly, the *d*_S_ distribution analysis across the *D. catalonica* paranome shows a canonical exponential decay, lacking the characteristic secondary peak indicative of a putative WGD (Vanneste et al. 2013) (supplementary fig. S1). Lastly, the phylogenetic relationship among *Dysdera* spiders with known genome sizes strongly supports that the genome size reduction (GSR) occurred only once, at the phylogenetic branch leading to the endemic species, thereby representing a derived evolutionary state. Specifically, the genome size of non-monophyletic mainland species ranges from 2.9 to 3.3 Gb, while those of Canarian species ranges from 1.5 to 1.7 Gb (Fig. 1a). Interestingly, our flow cytometry analyses indicate that *D. coiffaiti*, endemic to the Madeira islands and originated from an independent colonisation event from the mainland (Crespo, Silva, et al. 2021), shows a non-reduced genome (3.1 Gb).

### The GSR impact in chromosomes and repetitive elements

The genome of the Canarian species shrank by 1.73 Gb, representing a GSR of 52.7% (Fig. 2a and c, Table 1; supplementary table S5), mostly encompassing TE elements. The GSR is relatively consistent across autosomes, ranging from 50.0% to 55.4%, but notably lower (34.4%) for the X chromosome (supplementary tables S6a-c). This disparity is particularly pronounced in the non-repetitive fraction of the genome, with a GSR ranging from 34.2% to 39.6% for the autosomes, and only a 13.2% for the X chromosome (Table 2; supplementary table 6c). The reduction of specific TE orders and superfamilies further supports the contrasting evolutionary dynamics between the X chromosome and the autosomes, with the X chromosome consistently showing the lowest rates of contraction (supplementary table 6c).

DNA transposons represent the most abundant TE class in both species, occupying a similar proportion of the *D. catalonica* (42.0%) and *D. tilosensis* (39.3%) genomes. More specifically, the hAT and TcMar superfamilies are the largest contributors to genome size, together accounting for ∼30% of it. Unlike DNA transposons, retrotransposons show more pronounced differences between both species, encompassing 28.2% and 22.6% of the *D. catalonica* and *D. tilosensis* genomes, respectively (Fig. 2b; supplementary tables S5 and S6). Furthermore, there are important differences across specific retrotransposon orders and superfamilies, with superfamilies of the LTR order reduced up to ∼80% in *D. tilosensis* (supplementary table S5).

The distribution of DNA divergence among TE copies from the same family or superfamily differs substantially between species. We found that, globally, *D. tilosensis* exhibits higher Kimura divergence estimates, suggesting that its TEs are comparatively older than those in *D. catalonica* (Fig. 2d; supplementary fig. S2, supplementary table S7). Indeed, considering all REs, which are largely composed of TEs, the median divergence is 16.7% higher in *D. tilosensis* than in *D. catalonica*, a gap increasing to 20.7% for the X chromosome. Notably, this pattern differs between TEs classes, with retrotransposons, again, being the most differentiated class between mainland and insular species. For instance, the LTR order, which accounts for 60.1% of the retrotransposons in *D. catalonica* and only 26.7% in *D. tilosensis* (supplementary tables S5-7). In contrast, certain LINE superfamilies, such as RTEs and Dong-R4, exhibit lower divergence and occupy a larger fraction in the *D. tilosensis* genome. Nevertheless, the total base pairs contributed by these younger elements in *D. tilosensis* are minimal compared to all other TEs.

On the other hand, the GO enrichment analysis identified 147 biological processes (BP), 115 molecular functions (MF), and 25 cellular components (CC) significantly enriched among orthogroups lost in insular lineages (*p*-value < 0.01; supplementary fig. S3 and supplementary table S8). Notably, the most enriched GO terms include transposition and DNA integration (BP), DNA polymerase activity, DNA binding and transposase activity (MF), among others. Taken together, our findings indicate a change in transposition activity associated with the colonisation process, since most TEs exhibit limited (perhaps no activity) in insular species, except for certain LINE superfamilies characterised by young, potentially active elements.

### The impact of the GSR in genic and intergenic regions

We also examined how this genome reduction affected other genomic elements beyond REs, and found that gene lengths are significantly shorter in both insular species (Fig. 3a). However, the GSR had a minor effect on its coding sequences (CDS), with *D. tilosensis* showing only a 3.9% reduction in the length of 1:1 orthologous genes (7,547 cases). This reduction is even smaller, 0.2%, for genes located on the X chromosome (supplementary table S10, see also supplementary fig. S4a). Instead, this GSR is much more evident in other genome features evolving under lower functional constraints, such as introns (Fig. 3b) and intergenic regions (Fig. 3c; supplementary tables S11 and S12). Specifically, introns are ∼30.0% shorter in *D. tilosensis* (*p*-value 1.39·10^-240^, Mann-Whitney U test; supplementary table S11), for both autosomes and the X chromosome (Table 2; supplementary table S11). And yet, the average intron length remains nearly twice longer on the X chromosome than in the autosomes. Notably, small introns (less than 300bp) exhibit a modest 1.6% reduction (*p*-value 1.76·10^-3^, Fig. 3b; supplementary table S11), consistent with their higher functional constraint, as they can contain regulatory elements. As for the intergenic length, as defined between two adjacent 1:1 orthologous genes maintained in synteny between *D. catalonica* and *D. tilosensis*, the GSR is reduced by 51.5% (*p*-value 1.97·10^-271^, Wilcoxon test; Fig. 3c; supplementary table S12), with the X chromosome showing again a more modest reduction (40.4%). Remarkably, the longer the intergenic region, the larger its contraction, with contraction proportions reaching 78% for intergenic regions larger than 1 Mb (supplementary table S12).

**Fig 3.**
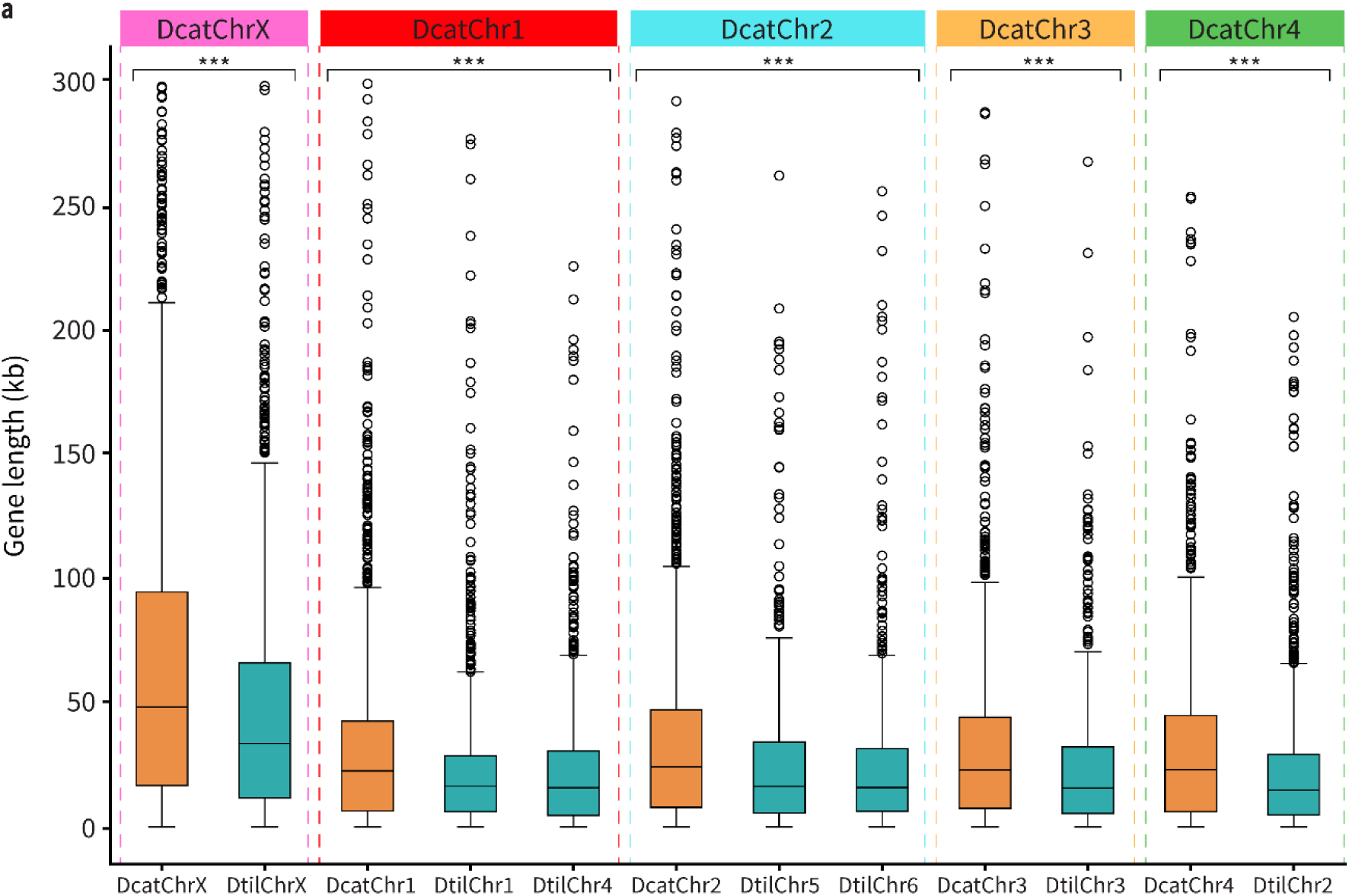
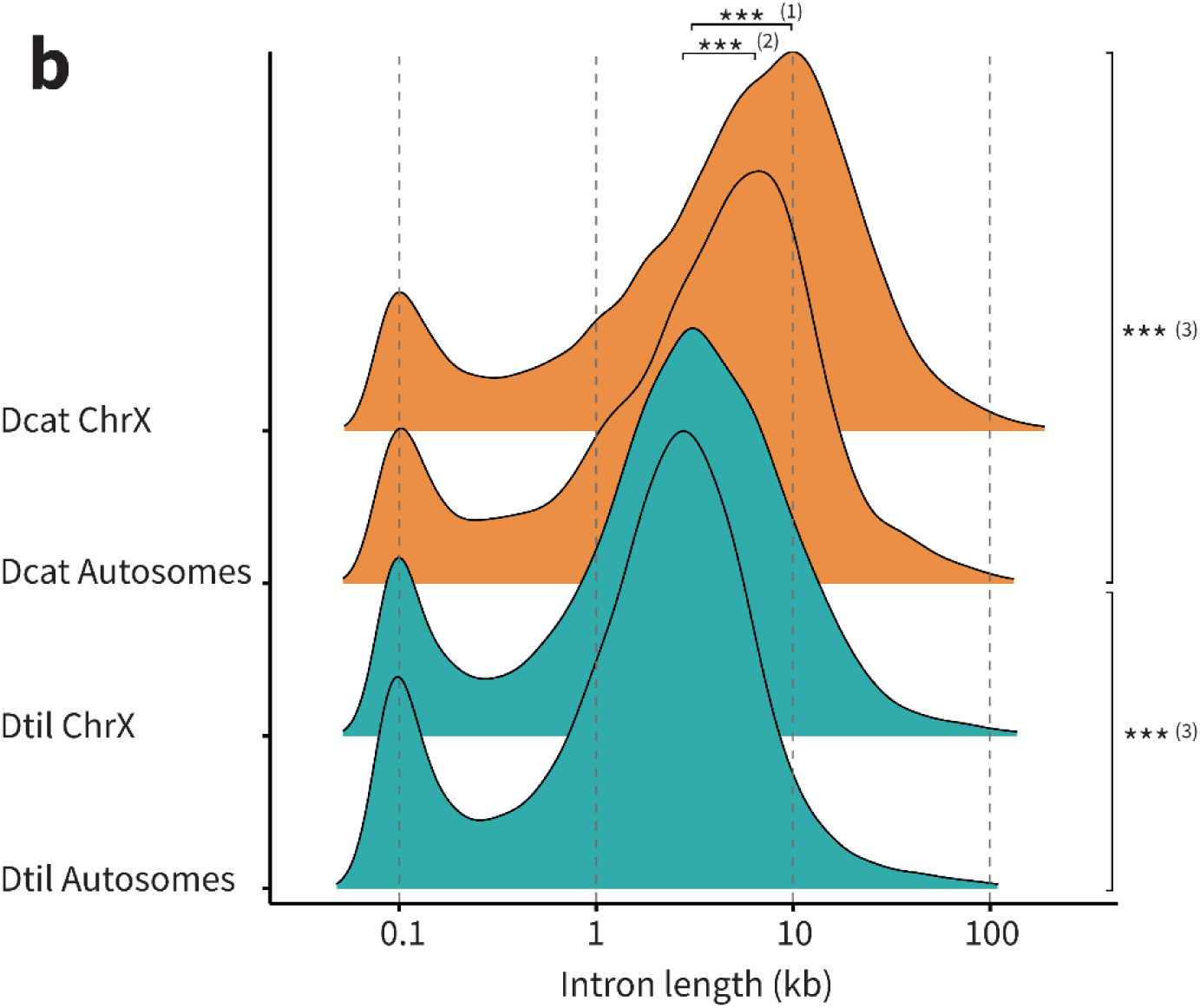
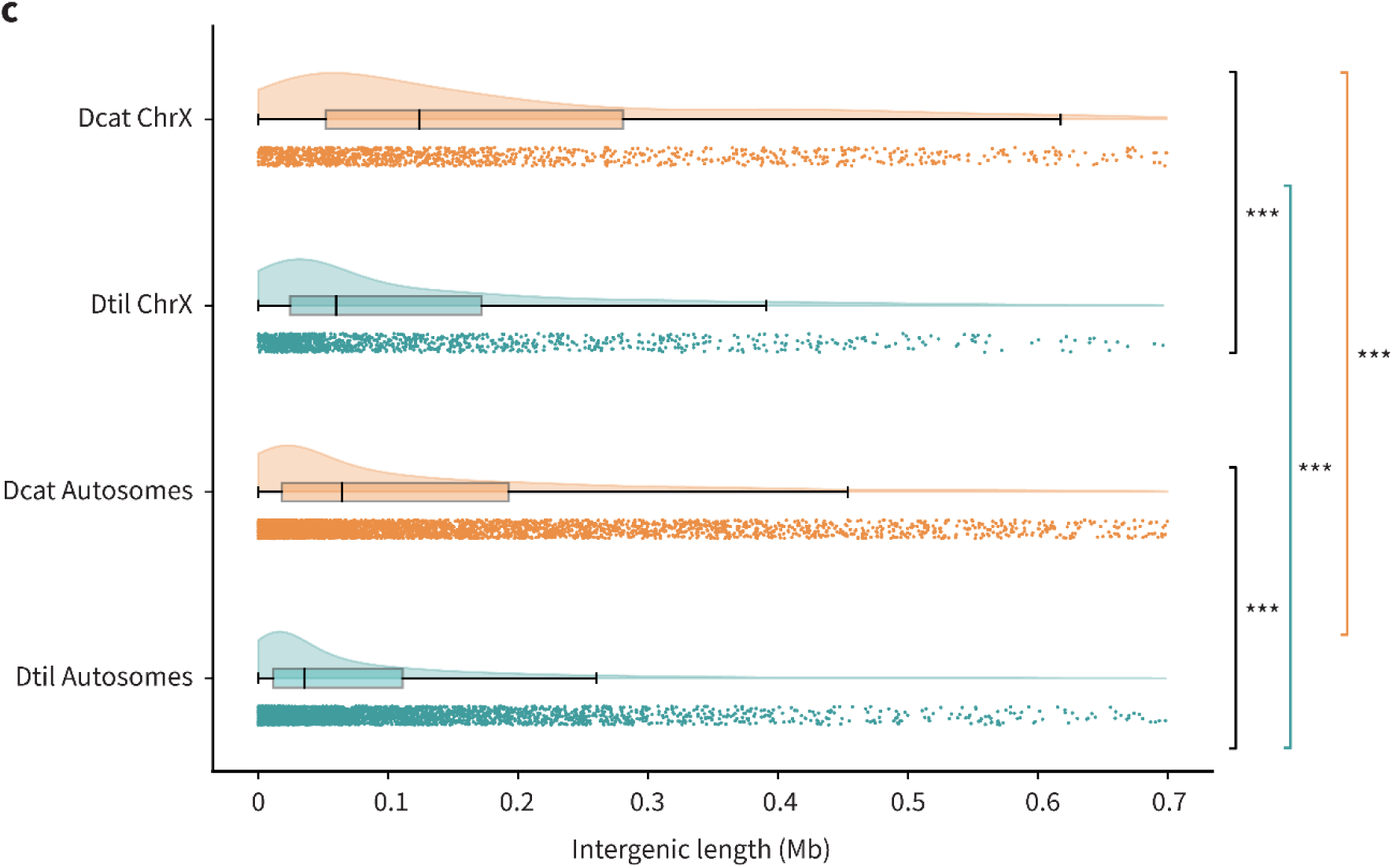
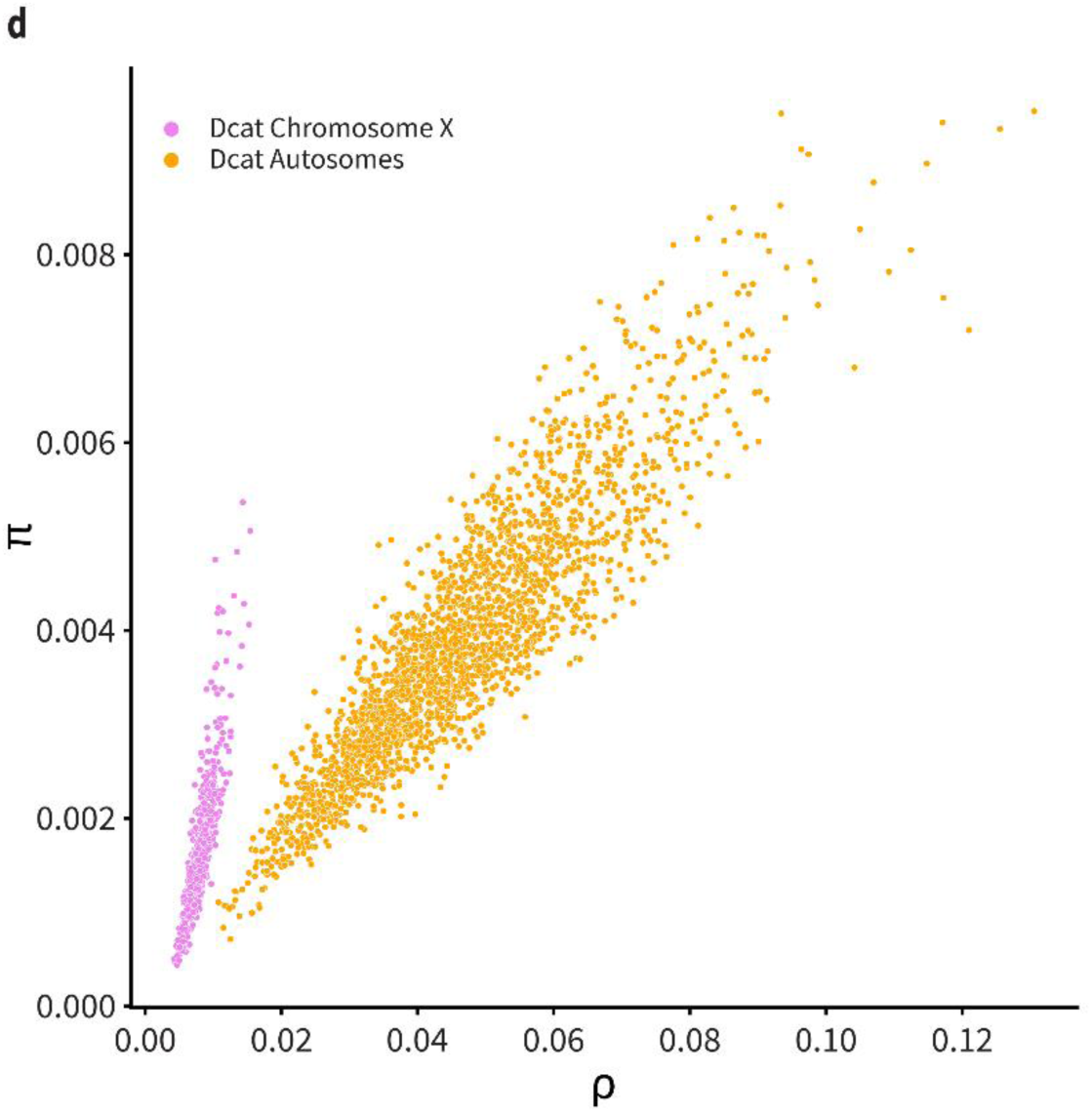
a) Gene length distribution in *D. catalonica* (orange) and in *D. tilosensis* (blue). Analysis based on 7,541 1:1 orthologous genes. Genes longer than 300 kb have been excluded for clarity. DcatChr1 gene lengths were compared to data from homologous chromosomes DtilChr1 and DtilChr4. Similarly, DcatChr2 gene lengths were compared to data from DtilChr5 and DtilChr6. Statistical significance was assessed with the Wilcoxon signed-rank test (*** *p*-value < 0.001). b) Intron length distribution in *D. catalonica* (orange) and in *D. tilosensis* (blue), based on 7547 1:1 orthologous genes. Both species show a bimodal distribution, with a peak of shorter introns at ∼100 bp. (1) and (2), indicate significant differences between the X chromosome and autosomes of *D. catalonica* and *D. tilosensis*, respectively, and (3) between X chromosomes or autosomes from the same species. Statistical significance was evaluated through the Mann-Whitney test (*** *p*-value < 0.001). c) Intergenic length distribution in *D. catalonica* (orange) and *D. tilosensis* (blue). Analysis based on 6,451 1:1 syntenic orthologous genes. Statistical significance between intergenic categories obtained with the Wilcoxon (*** *p*-value < 0.001). d) Correlation between π and ρ in 1 Mb genomic windows for *D. catalonica*. X chromosomes and autosomes are shown separately.

### Characterising the deletion bias in *Dysdera* species from the Canary Islands

Using available genome data from a mainland outgroup species, *D. scabricula* Simon, 1882, we estimated an average deletion and insertion lengths of 24 bp and 33 bp for introns, respectively, in the lineage leading to *D. tilosensis* (supplementary table S13 and methods Supplementary Material online). Nevertheless, the number of deletion events is higher than that of insertions (3.2 times), leading to a deletion bias (defined as deletion-to-insertion ratio) of 2.3 (supplementary table S13). This deletion bias in introns, however, cannot explain the observed GSR. Indeed, usingunpolarized data from local alignments (see methods in Supplementary Material online), we found that the vast majority (95.1%) of the genomic reduction owes to large, mainly intergenic, deletions. Specifically, we estimated that the average deletion and insertion lengths are 8.4 kb and 4.1 kb, respectively, with a deletion bias of 2.0, mirroring the overall genome contraction documented for insular species (supplementary table S14). In contrast to introns, where the number of events was also important, the deletion bias in intergenic regions was basically driven by indel length, with the X chromosome exhibiting lower deletion bias compared to the autosomes (supplementary tables S13 and S14).

### Analysis of functional and selective constraints

We first analysed whether weak selection could explain the observed GSR, using the G+C content and the effective number of codons as proxies for this selective force. We did not find any relevant differences between insular and continental species (supplementary figs. 4b and c, supplementary table S15). Their 1:1 orthologs genes show average G+C contents of 42.1% and 41.8% for *D. catalonica* and *D. tilosensis*, respectively, with similar proportions in autosomal and X-linked genes. Similarly, their average ENC’ values are 58.6 and 58.7 in *D. catalonica* and *D. tilosensis*, respectively, with minimal differences between autosomes and the X chromosome (supplementary table S15).

We also sought to identify shifts in selective constraints, either an intensification or a relaxation of the selective constraints during the diversification of the genus *Dysdera*, with special focus on the lineage that colonised the Canary Islands. We conducted this analysis based on the 4,689 1:1:1 orthologous genes shared between *D. catalonica, D. tilosensis* and *D. silvatica* (see Materials and Methods and supplementary table S16). After correcting for multiple testing, we identified 1,732 and 398 genes evolving either under intensified or relaxed selection, respectively (supplementary table S16). Notably, the branches leading to insular lineages show 4.75 times more genes undergoing intensification than relaxation (Models 1-5; supplementary table S16). In contrast, this ratio is balanced around 1.07 in the branch leading to *D. catalonica* (Model 6; supplementary table S16). These results show that selection was significantly stronger in the insular lineages. Interestingly, both synonymous (*d_S_*) and nonsynonymous (*d_N_*) substitution rates are approximately twice as high along the ancestral lineage leading to the two insular species (branch A, in supplementary table S17). This difference, however, is not statistically significant when assessed using tests of relative coding substitution rates.

### Levels of nucleotide polymorphism and recombination

To assess the impact of *N*_e_ on selection intensification, we analysed patterns and levels of nucleotide diversity and recombination. We consistently found greater levels of nucleotide diversity in *D. tilosensis* (and in *D. silvatica*) than in *D. catalonica*, in both autosomes (π = 0.016 vs. π = 0.0039), and the X chromosome (π = 0.009 vs. π = 0.0016) (supplementary table S18a). A similar pattern was observed for the recombination parameter, both in autosomes (ρ = 0.140 vs. ρ = 0.053), and in the comparison of the X chromosome between *D. silvatica* and *D. catalonica* (ρ = 0.025 vs. ρ = 0.008). As observed in other organisms (Ellegren and Galtier 2016), nucleotide diversity and recombination correlated significantly throughout the genomes of the the three species (τ = 0.79, τ = 0.19 and τ = 0.75, for *D. catalonica*, *D. tilosensis* and *D. silvatica*, respectively) (Fig. 3d; supplementary fig. S5 and supplementary tables S18b and c), supporting that the enhanced efficacy of selection in the Canarian species was ultimately driven by their larger *N*_e_.

## Discussion

Different hypotheses have been proposed to explain genome size evolution across species (Elliott and Gregory 2015; Wright 2017; Blommaert 2020; Galtier 2024), involving adaptive and non-adaptive mechanisms. Nevertheless, our understanding of the forces driving genome size evolution remains profoundly limited, largely due to inter-related confounding factors (Charlesworth and Barton 2004). Our biological model, the red devil spider genus *Dysdera* from the Canary Islands, offers a unique opportunity to investigate the key determinants of genome size variation. Our study benefits from newly generated chromosome-level assemblies to precisely determine the contribution of distinct genetic elements and events, circumventing the limitations of correlation-based studies based on distantly related species (Charlesworth and Barton 2004; Whitney et al. 2011; Galtier 2024). Therefore, our framework enables an accurate investigation of the patterns and mechanisms underlying genome size variation, as well as a critical evaluation of the most influential adaptive and non-adaptive hypotheses on the observed genome reduction.

### Unexpected genome size reduction in island-endemic species: the paradox of adaptive radiations

At first glance, the twofold difference in genome size between insular and mainland species could be easily misinterpreted as a WGD event, consistent with several studies reporting multiple rounds of WGD in chelicerates (Shingate et al. 2020), but see Thomas et al. (2024). However, we provide clear-cut evidence for a GSR in the lineage leading to island species. Nevertheless, it remains unknown whether this reduction predated the colonisation event (Bellvert, Pollock, et al. 2024), or was a direct consequence of it. Moreover, the observation of shorter genomes in endemic species of small oceanic islands is quite unexpected. In fact, the founder effects and bottlenecks associated with colonisation are expected to strengthen genetic drift, reduce genetic diversity and weaken the efficiency of purifying selection in removing transposons, and consequently leading to an increased genome size (James et al. 2016; Cerca et al. 2023; Yang et al. 2024). That all flow cytometry results (Fig. 1a) point to similar genome size for all Canarian species (1.7 Gb) and for all mainland relatives (3 Gb) suggests that genome size is in equilibrium in both groups of species. Our study sheds light on this paradox, suggesting an important role of factors rarely accounted for in island colonisation models, such as the increase in the effective population size, recombination and mutation rates.

### The genome reduction in Canary Island-endemic species is not adaptive

The remarkable GSR observed in the genus *Dysdera* is difficult to reconcile with the most popular adaptive hypotheses. First, there are no apparent differences in body size or life-history traits between species with large and small genomes. Unfortunately, as is common in non-model organisms, there is limited data on key traits—such as metabolic rate, cell size (e.g., egg or clutch size), and developmental time—have not been systematically investigated across the focal species. Both groups live in a variety of habitats and are able to feed on woodlice with the same range of preferences (Řezáč et al. 2018; Bellvert et al. 2023). Thus, it is unlikely that genome size, through its association with cell size, was directly targeted by selection in the insular species (Gregory and Hebert 1999; Cavalier-Smith 2005; Tsukaya 2013). Second, the observed change in genome size is also very difficult to explain under the genome streamlining hypothesis (Hessen et al. 2010; Stelzer et al. 2023), which proposes phosphorus and/or nitrogen limitation as the main driver of GSR. Studies in oceanic islands suggest that these limitations would only appear in young (N) or very old (P) islands (Vitousek and Farrington 1997). Furthermore, *D. coiffaiti*, a species endemic from Madeira Islands, a Macaronesian archipelago geologically similar in age to the Canary Islands (van den Bogaard 2013; Ramalho et al. 2015), has a genome size similar to that of the mainland species (Fig. 1a). Although *Dysdera* likely colonised the Canary Islands early after their formation (Bellvert, Dimitrov, et al. 2024), reports of both reduced and not reduced genomes in Macaronesian plants (Suda et al. 2005; Pellicer and Fernández 2023) indicate that the availability of essential nutrients has not been a limiting factor in the latter archipelago. This undermines the genome streamlining hypothesis as an explanation for the striking differences in genome size among *Dysdera* species.

### A genome-wide large-segmental deletion bias explains the observed genome reduction

Our analysis reveals that the reduction of repetitive sequences, particularly TEs, explain most of the GSR (supplementary table S5). Moreover, we found that this reduction is relatively consistent across autosomes and comparatively smaller in the X chromosome. That this reduction is dominated by TEs is unsurprising, as these elements are predominant components of most animal genomes (Marino et al. 2024). However, the GSR also extends to other genomic features, especially if evolving under low functional constraint, such as intergenic regions or paralogous copies (supplementary tables S4, S5 and S12). The main molecular mechanism involves large intergenic deletions (segmental deletions), as small indels contribute much less. All these findings suggest that ectopic recombination, fuelled by TE sequences, is the main mechanism for DNA loss (Montgomery et al. 1991). Indeed, the high fraction of interspersed REs provides a suitable substrate for unequal crossing over resulting in large deletions (Balachandran et al. 2022). Therefore, unlike other studies (Gregory 2004; Laurie et al. 2012), we can rule out the mutational equilibrium hypothesis (Petrov 2002) in explaining the observed GSR. The observed deletion bias, driven by large deletions, closely aligns with the accordion model proposed by Kapusta et al. (2017) in birds. Contrary to our observations, however, this hypothesis places little emphasis on TE mobilisation but highlights that variations in life-history traits, such as metabolic requirements, determine DNA loss rates.

### Long-term effective population size drove GSR in the endemic *Dysdera* spiders

Naively, our results seem to contradict the expectations underlying one of the most influential hypotheses explaining genome size variation, the mutational hazard hypothesis (Lynch 2011; Smith 2016: 201; Lynch et al. 2023). Indeed, the smaller *N*_e_ expected on oceanic islands would reduce the efficacy of selection (i.e., genome-wide relaxation of selective constraints). This, in turn, would lead to an increase in genome size due to the accumulation of deleterious DNA that cannot be efficiently purged by purifying selection. However, our results suggest that the insular species experienced an increase in the long-term *N*_e_, accompanied by an intensification of natural selection following their colonisation of the Canary Islands. According to the mutational hazard hypothesis, we would actually expect the observed reduction in genome size. Furthermore, nucleotide polymorphism and recombination rates are higher in the insular genomes and correlate in all surveyed species (Fig. 3d; supplementary table S18). Since both parameters depend on the *N*_e_, and the evolutionary rates following the split between *D. catalonica* and the lineage leading to the insular species are not significantly different (RRT, *p*-value > 0.05 considering both coding or nucleotide-based divergence), our results suggest that *N*_e_, rather than the mutation rate, is the key factor in determining the observed differences in genome size. Our findings align with previous studies indicating that census size, at least on islands, may be a poor indicator of long-term effective population size (James et al. 2016; Escuer et al. 2024). Nevertheless, although the observed differences in *d_s_* are directionally consistent with a potential increase of the mutation rate in the internal branch ancestral to the island species clade, encompassing the colonization event, they are not statistically significant, suggesting that mutation rate is unlikely to be the primary driver of the observed GSR.

The greater GSR in the autosomes compared to the X chromosome also supports the mutational hazard hypothesis. The *N*_e_ is expected to be lower in X chromosome than in autosomes (a reduction of 0.75 (Lynch and Conery 2003; Luiselli et al. 2024); assuming equal number of males and females), resulting in a weaker intensification of natural selection acting on this chromosome. Although changes in *N*_e_ should theoretically affect both X chromosome and autosomes proportionally – resulting in a similar selection intensification on both– several potential confounding factors could decouple this relationship. These include sex-ratio imbalances, male hemizygosity, sexually antagonistic mutations, dosage compensation and the so-called faster-X effect (Vicoso and Charlesworth 2006; Charlesworth 2009). Nonetheless, the greater GSR observed in autosomes may be more easily explained by the comparatively reduced effective recombination rate of the X chromosome, which is limited to females due to male hemizygosity (Nam and Ellegren 2012; Charlesworth et al. 2018; Moran et al. 2018).

In summary, our findings are compatible with a non-adaptive genome size reduction, with purifying selection as the primary driver. We therefore propose the hypothesis that the GSR was driven by an increase in the long-term *N*_e,_ which enhanced the efficiency of selection. This process may have been further reinforced by an increased effective recombination rate (Nam and Ellegren 2012). Therefore, the observed GSR is likely driven by the intensification of selection against mutations with slightly deleterious effects, such as those DNA insertions arising from transposition and gene duplications. Supporting this idea, we detected older TEs in the two island species, suggesting TE inactivation or reduced activity in their reduced genomes (Mérel et al. 2024). This evidence aligns with the observed enrichment of GO terms related to DNA transposition and integration associated with genes lost in the island species (supplementary fig. S3 and supplementary table S8). Therefore, the most plausible scenario explaining the GSR, is the result of both, a reduced rate of DNA gain and an increased occurrence of large deletions. Specifically, DNA gains driven by TEs may be constrained by intensified selection and reduced insertion rates due to diminished mobilization. Additionally, the deletion bias may have increased concurrently due to enhanced effects of the ectopic recombination effects associated with the higher recombination rate. Together, these processes may have contributed to establish a new genome size equilibrium in the Canary Island species.

Our study revealed a remarkable and unexpected result: an increase in the efficacy of selection in small oceanic islands species that have undergone a founder (probably involving a bottleneck) effect (Cerca et al. 2023). However, the impact of genetic drift on island colonisation events remains poorly understood (James et al. 2016; Yang et al. 2024). Given that the colonisation of the Canary Islands occurred approximately 25 Mya (Crespo, Isamberto, et al. 2021), sufficient time has passed for the genomic signatures of the colonization event to be eroded. On the other hand, the more stable climatic conditions on the islands compared to the mainland, along with potentially larger-than-expected census sizes, may also have contributed to a high long-term historical *N*_e_ in the insular species of the genus *Dysdera.* In fact, this scenario, which is consistent with Escuer et al. (2024) results, would account for the intensified selection observed in these lineages. Globally, our findings provide strong support for the non-adaptive mutational hazard hypothesis as the primary explanation for genome size evolution in Canary Islands *Dysdera.* Given that our dataset includes only three species —only one of which is from the mainland, —and that few studies have compared life-history traits between mainland and island species, additional genome surveys, coupled with more systematic comparisons of life-history traits, are necessary to strengthen and corroborate this hypothesis. This will definitely contribute to a deeper understanding of the forces shaping species diversification in oceanic islands and the biological significance of genome size evolution.

## Material and Methods

### Study design, sampling, DNA extraction, sequencing and genome assembly

This study relies on chromosome-level assemblies of three spider species belonging to the genus *Dysdera*: two endemics to the Canary Islands, *D. silvatica* and *D. tilosensis*, and one from the European mainland, *D. catalonica* (Fig. 1b). Specimens of *D. tilosensis* (Fig. 1c) were sampled in Gran Canaria island (Canary Islands, Spain) in 2015 and 2022, while those of *D. catalonica* in the Parc Natural del Montseny (Catalonia, Spain) in 2020 (supplementary table S1). The genome of *D. silvatica* was obtained from public databases (Escuer et al. 2022), whereas those for *D. tilosensis* and *D. catalonica* were assembled at chromosome level in this study (Fig. 1d). We estimated genome sizes of several islands and mainland *Dysdera* species (Fig. 1a) using flow cytometry as in Sánchez-Herrero et al. (2019). The genome size of *D. crocata* had been previously determined (Gregory and Shorthouse 2003).

DNA extraction was performed using the Gentra Puregene Cell kit (Qiagen) for *D. tilosensis* and *D. silvatica* (for individual resequencing), and the Blood & Cell Culture DNA Mini Kit (Quiagen) for *D. catalonica*. RNA extraction for the annotation of *D. catalonica* genome was performed as in Vizueta et al. (2017) and in Frías-López et al. (2015). To obtain the chromosome-level assemblies, we sequenced the genomic DNA with PacBio CLR technology (with 101X and 57X coverage for *D. catalonica* and *D. tilosensis*, respectively), and performed the scaffolding using the Omni-C technology (Dovetail Genomics) (supplementary table S2). mRNA sequencing (66 Gbp in *D. catalonica*) and whole-genome individual resequencing (with a sequencing depth of 39X, 37X and 41X for *D. catalonica*, *D. tilosensis* and *D. silvatica*, respectively), were performed using 150bp paired-end reads on the illumina NovaSeq 6000 platform (supplementary table S2). Genome assembly was performed in two steps: an initial assembly with wtdbg2 v2.5(Ruan and Li 2020), followed by a scaffolding step with HiRise (Putnam et al. 2016). Assembly completeness was determined with the BUSCO-v2.5.4 pipeline (Seppey et al. 2019), using arachnida (odb10; 2,934 genes), arthropoda (odb10; 1013 genes) and eukaryota (odb10; 255 genes) datasets (Kriventseva et al. 2019).

### Annotation of repetitive elements

RepeatModeler-v2.0.3(Flynn et al. 2020) was employed for *de novo* identification of REs, using the LTR discovery pipeline (-LTRStruct). Sequences categorised as “unknown” were then re-classified with DeepTE (Yan et al. 2020) using the metazoan model (-sp M). The RE libraries obtained for the three species were merged into a single database, which was used as input for masking with RepeatMasker-v4.1.2 (http://repeatmasker.org), configured with Dfam 3.5 (Storer et al. 2021) and RepBase-20181026 (Bao et al. 2015), and run using sensitive search (-s). The TEs identified by RepeatMasker were classified using custom Python scripts based on TEs class, subclass, order and superfamily, following Wicker et al. (2007) and Wells and Feschotte (2020).

### Structural and functional genome annotation

Gene models were generated with BRAKER-v2.6.4 in ETPmode, using protein and RNAseq evidence (Stanke et al. 2006; Stanke et al. 2008; Hoff et al. 2019; Brůna et al. 2021). Protein evidence included orthologous sequences from five species of the *Dysdera* genus (including *D. silvatica* and *D. tilosensis*) (Vizueta et al. 2019), jointly with those from arthropoda_odb10 from OrthoDB v10 (Kriventseva et al. 2019). The *D. tilosensis* and *D. silvatica* RNAseq data were obtained from public databases (Vizueta et al. 2017). RNAseq reads were trimmed with Trimmomatic-v0.39 (Bolger et al. 2014) using the following parameters (LEADING:3, TRAILING:3, SLIDINGWINDOW:4:15, MINLEN:50) and mapped to their respective reference genomes with STAR-v2.7.10a (Dobin et al. 2013).

Functional annotation of the predicted gene-models was performed by running BLASTP-v2.13 searches (E-value = 1·10^-3^) against the SwissProt database (downloaded in August 2022), and a custom arthropoda database (Vizueta et al. 2017; Escuer et al. 2022). To further improve functional annotations, we additionally applied InterProscan-v5.57-90.0 with default parameters (Jones et al. 2014) and EggNOG-mapper-v2.1.9 (Huerta-Cepas et al. 2017; Cantalapiedra et al. 2021) in ultra-sensitive mode. Finally, we used the accessory BRAKER2 script selectSupportedSubsets.py to classify predicted genes into those fully supported by external evidence (SBS genes) and the remaining genes.

### Synteny, orthology and analyses of insertion-deletion patterns

To elucidate the putative role of WGD in genome size evolution, we analysed the synteny across species. We conducted pairwise genome alignments between the three species using minimap2-2.24 (Li 2018) with the preset “-x asm20”, which maps genomes with up to 10% sequence divergence. Results were visualised using Dgenies-1.2.0 (Cabanettes and Klopp 2018) and circlize-0.4.15 package (Gu et al. 2014). We also used GENESPACE-v1.3.1 (Lovell et al. 2022), which integrates evidence from gene collinearity and sequence similarity, to infer syntenic blocks. We examined the paranome’s *K*_S_ distribution (Blanc and Wolfe 2004), using the wgd-1.1.2 tool (Zwaenepoel and Van de Peer 2019). The ancestral genome size state was determined from flow cytometry data, in a phylogenetic context and assuming a parsimonious criterion.

Orthogroups across the three *Dysdera* species were determined using Broccoli-v1.2 (Derelle et al. 2020) and OrthoFinder-v2.5.4 (Emms and Kelly 2019), applying two consecutive rounds of refinement. Briefly, Broccoli was initially used to define the primary orthogroups. Genes unclustered by Broccoli were subsequently grouped into additional orthogroups using OrthoFinder. This combined approach yields four orthogroup classes: 1:1 and N:N orthogroups from Broccoli, and N:N or singletons groups resulting from the subsequent OrthoFinder analysis.

We estimated the number and length of different genomic features from the annotation files of each species, including genes, CDS, introns and intergenic regions. These analyses used the data obtained from genome annotation GFF3 files, jointly with 1:1 orthogroup information. However, to exclude potentially misleading orthogroups, we previously conducted several filtering steps, using custom Python scripts and BEDTools-v2.31.0 (Quinlan and Hall 2010). Statistical significance for paired and non-paired data was assessed through Wilcoxon signed-rank and Mann-Whitney U tests, respectively, using R version 4.12 programming language (R Core Team, 2021).

We estimated the patterns of insertion and deletion events (and deletion bias), applying two distinct, complementary approaches (see methods in Supplementary Material online). In the first one, we identified indels in introns well-delimited by orthologous flanking exons, previously aligned using MAFFT-v7.31 (Katoh and Standley 2013). To polarise evolutionary events, we incorporated genomic information from an outgroup species (*D. scabricula;* kindly provided by Miquel A. Arnedo research group), which consist in low-coverage sequencing using data (short-reads), as well as published genome data from the endemic species *D. silvatica* (Escuer et al. 2022). Given the high sensitivity of this analysis to alignment quality, we restricted the analysis to introns from 1:1 orthologous genes. Second, we estimated indels along the whole genome using information of local aligned regions between *D. catalonica* and *D. tilosensis* obtained with minimap2 (Li 2018).

### Recombination and nucleotide diversity analysis

We estimated the per-site nucleotide diversity (π) using VCFtools-v0.1.16 (Danecek et al. 2011) and population-scaled recombination rate (ρ) using the iSMC-v0.0.25 method (Barroso et al. 2019) with Illumina resequencing data from a single individual per species (supplementary tables S1 and S2, see detailed protocol in Supplementary Material online). Under the neutral model, the expected value of π is θ = 4×Ne×u (being u, the per-site per-generation mutation rate), and ρ = 4×Ne×*r*, (being *r*, the rate of crossover events per-nucleotide and per-generation). We also examined the correlations between π and ρ, separately for autosomes and the X chromosome, in 1Mb windows, and in well-characterised genomic regions, including intergenic and intronic regions (see Supplementary Material online). The statistical significance of the correlation between these two parameters was assessed with the non-parametric Kendall rank correlation coefficient (τ) using Python SciPy (Virtanen et al. 2020).

### Functional constraints in protein-coding genes

We tested for changes in the selective constraints of protein-coding genes during the diversification of the surveyed species, and after the split of the island endemic and mainland ones (Fig. 1a). We applied two complementary approaches on the 1:1 orthologs dataset (as obtained by Broccoli). First, we estimated the G+C content using the GC function in Biopython-v1.78 (Cock et al. 2009), and codon usage bias as the effective number of codons (ENC’) (Novembre 2002), using coRdon-v1.12.0 R package (Elek et al. 2019). Second, we assessed shifts in selective constraints on protein-coding genes using the method in RELAX, which is implemented in the HyPhy-v2.5.8 package (Pond et al. 2005; Wertheim et al. 2015). For this analysis, we compared six different branch models, each accounting for changes in selective constraints in specific lineages. We also estimated lineage-specific *d*_n_/*d*_s_ ratios using the codeml program from the PAML-v4.10.7 software (Yang 2007). We also used the relative rate test (RRT) implemented in Hyphy to compare the evolutionary rates after the split of the lineages leading to *D. catalonica* and the island endemic species. For that we built a concatenated CDS alignment with information from 1:1:1 orthologs among *D. catalonica, D. tilosensis* and *D. silvatica* (∼7 million CDS sites), and analysed the data using the model MG94W9 and GRM, to test coding- and nucleotide-based evolutionary rates, respectively. In both cases, parameters shared by all branches and branch lengths are estimated independently. We used *D. scabricula* as the outgroup species, *D. silvatica* or *D. tilosensis* as the ingroup 1, and *D. catalonica* as the ingroup 2.

### GO enrichment analysis

We obtained the GO terms associated with the genes annotated in the three species using InterProScan v.5.57–90.0 (Jones et al. 2014). The GO enrichment analysis was performed using the R packages GSEABase v.1.60.0 (Geistlinger et al. 2021; Morgan et al. 2024) and GOstats v.2.64.0 (Morgan et al. 2007), selecting as candidates those genes that belong to the orthology group N:0:0 (i.e., genes present in *D. catalonica* but absent in *D. tilosensis* and *D. silvatica*). In this analysis, we only include those genes supported by external evidence (SBS genes). We ran GOstats with the conditional option, and considered GO terms to be significantly overrepresented if their associated *p*-value were below 0.01. We also used the R packages clusterProfiler v.4.8.1 (Wu et al. 2021) and AnnotationForge v.1.42.0 (Carlson and Pagès 2024) to transform the GOstats results into an enrichResult object and plotted them using the R packages enrichplot v.1.20.0 (Yu) and ggplot2 v.3.4.2 (Wickham 2016).

## Supporting information

Supp. Meterial

## Data, Materials, and Software Availability

The whole-genome shotgun projects have been deposited at DDBJ/ENA/GenBank under accession numbers XXXXX, XXXXX and Bioproject ID PRJNA1270787. The Omni-C, PacBio CLR, WGS Illumina and mRNA sequencing data are also included in the Bioproject repository (SRR33784497 - SRR33784507) Structural and functional annotations are available in https://github.com/molevol-ub/Dysdera_CatTil_genomes.

## Acknowledgements

This work was supported by Ministerio de Ciencia e Innovación (MCIN/AEI/10.13039/501100011033) grants (PID2019-103947GB, PID2019-105794GB, PID2022-137758NB-I00, PID2022-138477NB, and an FPI fellowship to V.A.P. PRE2020-095592). Additional funding was provided through a predoctoral fellowship from the Universitat de Barcelona to M.O-M. (PREDOCS-UB 2021), and by Comissió Interdepartamental de Recerca i Innovació Tecnològica (2021SGR00279, 2021SGR0689). We gratefully acknowledge the Cabildo of Gran Canaria and the Canarian Government for granting collection permits and providing support with accommodation and logistics during fieldwork. We also thank Silvia Adrián-Serrano, Adrià Bellvert and Alba Enguídanos for sharing karyotype data, low-coverage whole-genome sequencing information for *D. scabricula*, and samples of *D. catalonica*.

