## Supplementary material for "How did evolution halve genome size during an oceanic island colonization?": Supp. Figures

**SUPPLEMENTARY FIGURES**

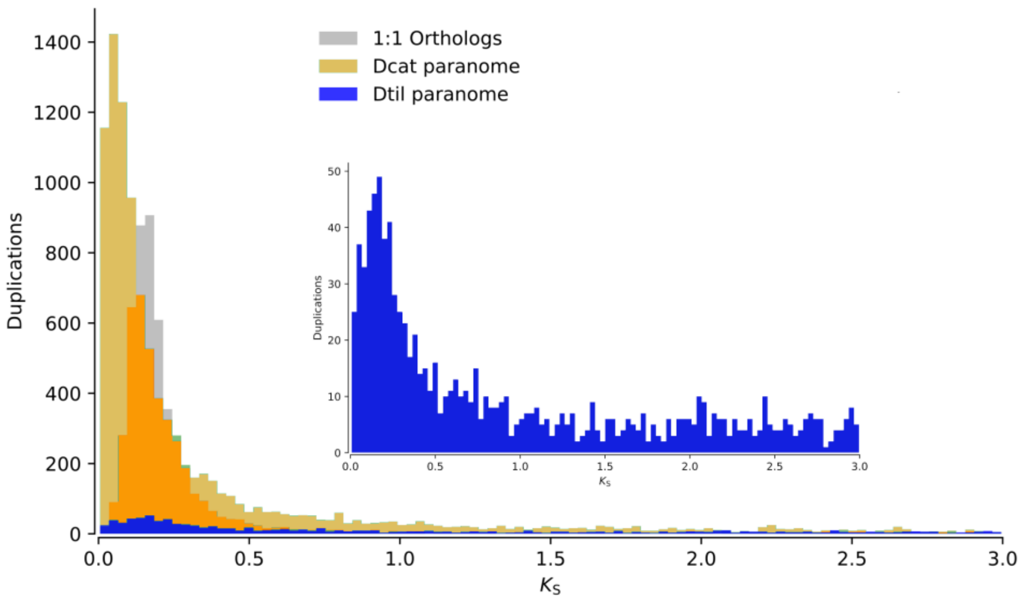


Fig. S1. Distribution of the synonymous nucleotide divergence ($k_s$) between orthologous and paralogous genes in *D. catalonica* (orange) and *D. tilosensis* (blue). The upper-right-panel shows a zoom of the paranome in *D. tilosensis*. The lack of secondary peaks in the $k_s$ distributions excludes explosive episodes of gene duplication.

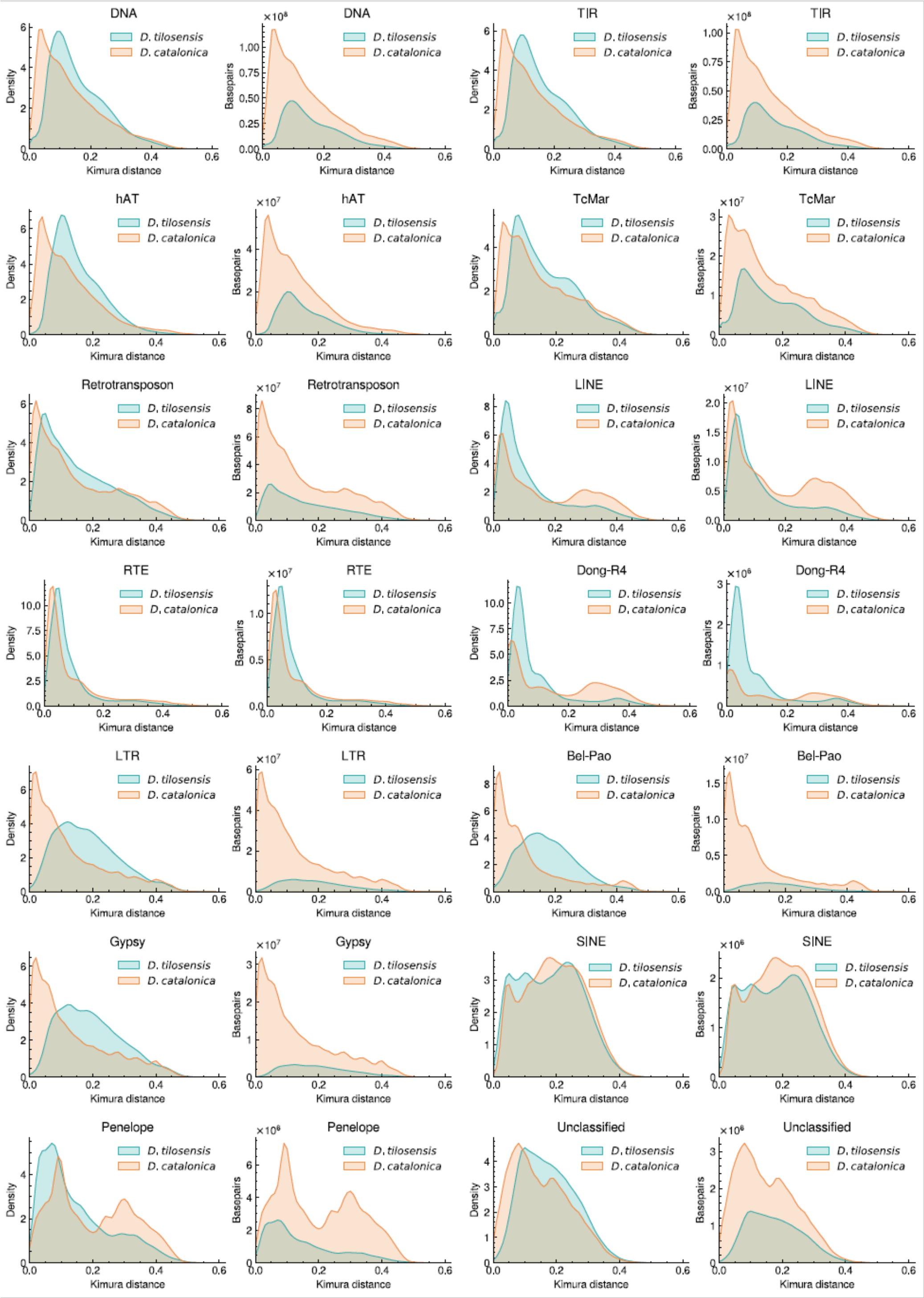


Fig. S2. Kernel density estimates for Kimura divergence of different TE groups. The Kimura genetic distances are adjusted for the CpG sites.

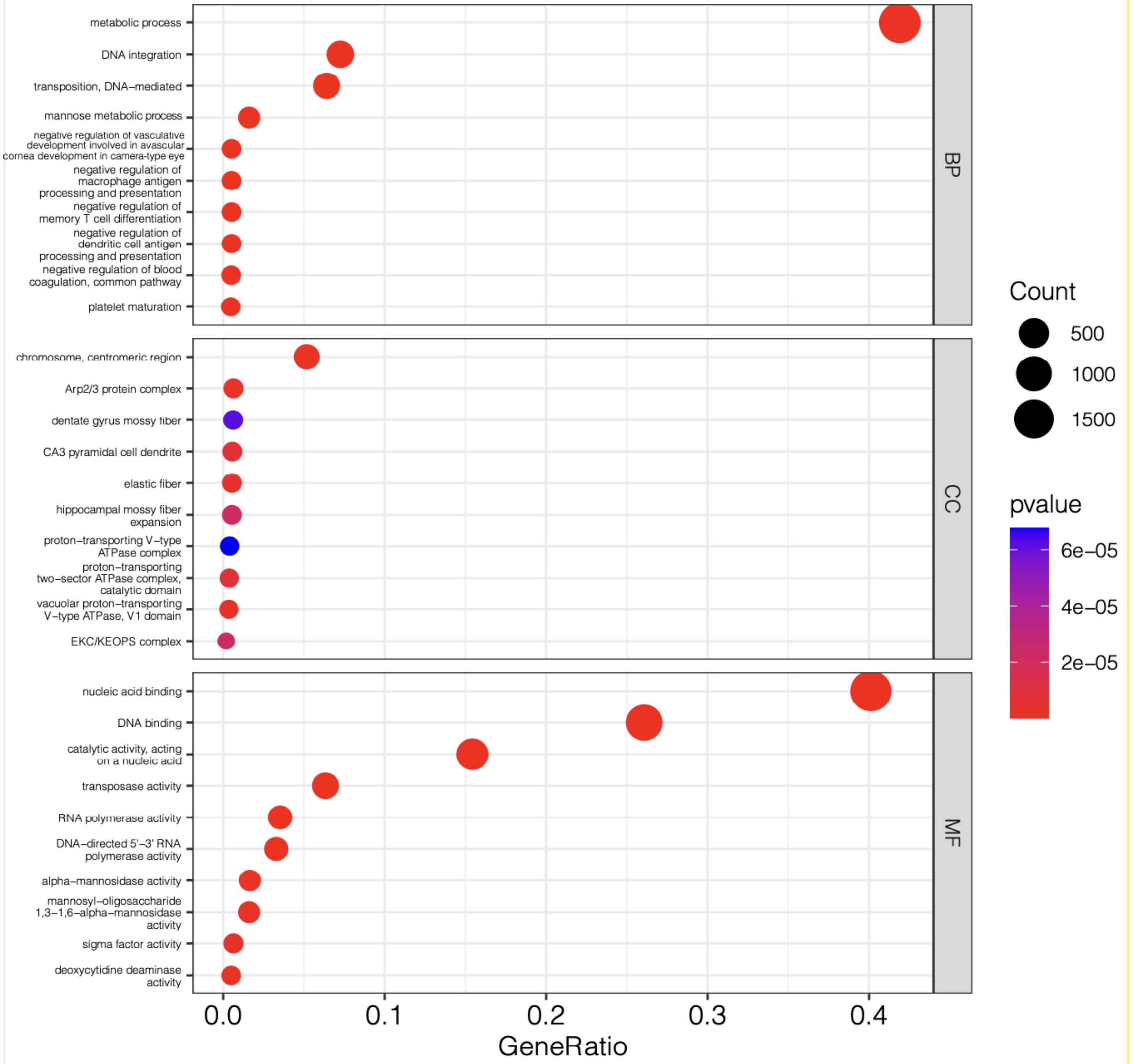


Fig. S3. Top 10 most significant GO terms in each GO category: Biological Process (BP), Cellular Component (CC) and Molecular Function (MF). Count refers to the number of genes in each GO term. GeneRatio refers to the fraction of the number of genes of interest associated with a specific GO and the number of total genes associated with that particular GO.

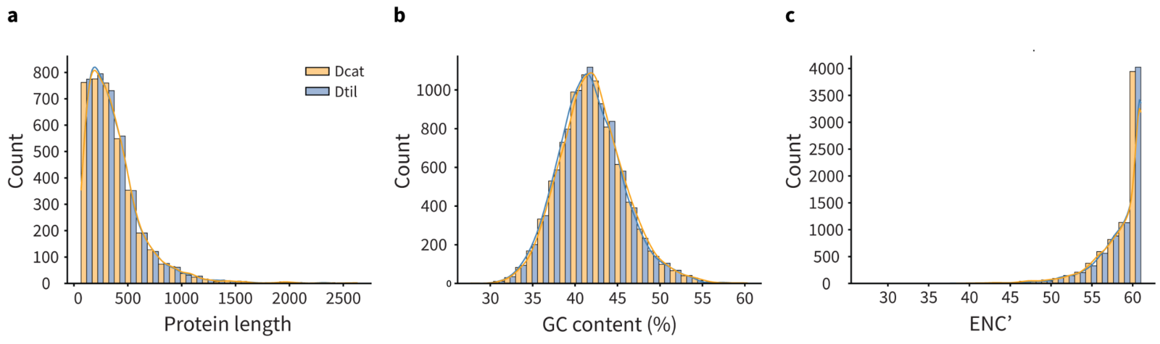


Fig. S4. Protein length, GC content and ENC' distribution between *D. catalonica* (orange) and *D. tilosensis* (blue). a) Analysis based on 3,784 proteins (1:1 orthologous genes encoding the same number of exons). b-c) Analysis based on 7,547 1:1 coding regions of orthologous genes.

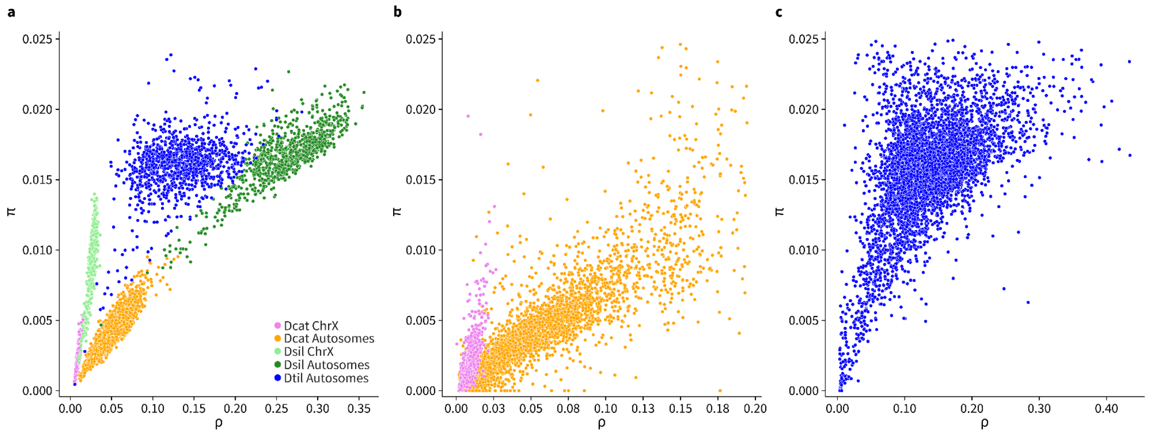


Fig. S5. Correlation between π and ρ. π values higher than 0.025 are not shown. a) Analysis in 1 Mb genomic windows for *D. catalonica*, *D. tilosensis* and *D. silvatica.* b-c) Analisis in syntenic intergenic regions (6,261 genes) in *D. catalonica* (panel b), and *D. tilosensis* (panel c). X chromosomes and autosomes are shown separately, except for *D. tilosensis* that there is no data from the X chromosome.
