## Supplementary material for "How did evolution halve genome size during an oceanic island colonization?": Supp. Meterial

### Supplementary Methods

**Analyses of Indels**

We determined the number and length of indel events applying two approaches. First, using information of well-characterized intron alignments of *D. catalonica* and *D. tilosensis* obtained with MAFFT v7.310 (Katoh and Standley, 2013), incorporating low-coverage short-read sequence data from an outgroup species (*D. scabricula*) to polarize indel events. Since low-coverage short-read sequence data precludes performing a fine analysis at high divergent regions, we performed a complementary analysis to analyse the patterns of insertions/deletions across larger chromosomal segments (mostly intergenic regions) from global alignment data *D. catalonica* and *D. tilosensis* obtained with minimap2-2.24 (Li, 2018). This second approach was performed, therefore, without polarying data.

**1. Assessing the indels patterns in intronic data**

We used data of the 4,981 1:1:1 orthologous genes between *D. tilosensis* (Dtil), *D. silvatica* (Dsil) and *D. catalonica* (Dcat) to infer the pattern (i. e., number and length) of indels in intronic regions. We used this conservative set of genes to ensure a reliable orthologous intronic relationship between insular and mainland species. To determine the indel pattern in *D. tilosensis* and *D. catalonica*, we polarize the insertions and deletions using low-coverage short-read sequence data from *D. scabricula* (Dsca).

*1.1. Outgroup sequence preparation*

Since only low-coverage Illumina sequence data are available on *D. scabricula,* we used a mapping strategy (against the *D. catalonica* reference sequence) to retrieve their CDS (hereafter, Dsca_1). For that, we used BWA-mem v0.7.17 (Li, 2013) to map the *D. scabricula* reads, Samtools v.1.14 (Danecek et al. 2021; Li, et al. 2009) to skip the alignments with mapping quality below 20 and BCFtools v1.16 (Li, 2011) to perform the variant calling, retaining only positions with high base calling quality (QUAL>=20), and to generate a consensus CDS sequences. Finally, these *D. scabricula* CDS' were translated to protein using a custom BioPython script (Figure SM1).

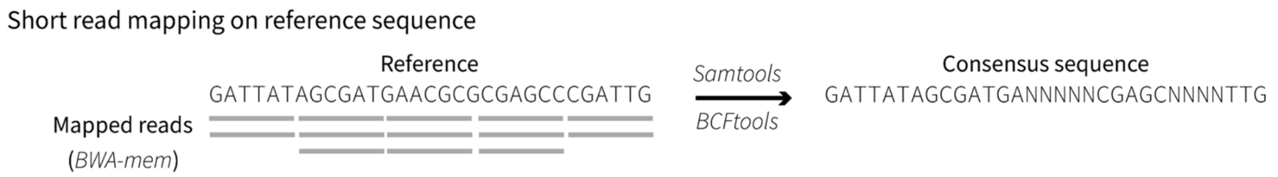


Figure SM1. Short reads (from *D. scabricula*) mapped against the reference sequence (*D. catalonica*).

#### *1.2 Protein alignment*

We first aligned the protein sequence data of *D. tilosensis* and *D. silvatica* using mafft `(--auto)`, and after that, we added *D. catalonica* and *D. scabricula* protein sequences with mafft `(--add)` option (Figure SM2a).

#### *1.3 Identify homologous exons*

We inferred matching pairs of adjacent exons using *D. catalonica* as reference. For that, we used a custom Python script to extract exon coordinates from GFF3 files, to associate them with their corresponding coordinates in the protein MSA (Figure SM2b-d). The resulting connections (pairs of matching exons) between pairs of species (*D. tilosensis* against *D. catalonica* and *D. scabricula* against *D. catalonica*) are represented as a graph (Figure SM2e), created with networkx-v3.1 package (Hagberg et al. 2008). Each node of the graph corresponds to one exon, and each edge to the alignment score between two homologous exons, ranging from 0 (no alignment) to 1 (complete alignment). We retained only those pairs of adjacent exons shared and perfectly aligned between *D. tilosensis, D. catalonica* and *D. scabricula* species. Exons with secondary connections are excluded since they can represent either misalignments or cases of fused exons (Figure SM2f).

#### *1.4. Create concatenates of CDS and introns*

For each pair of matching exons, we created a concatenate sequence that included the two adjacent exons and their delimited intron. For that, the DNA sequences of *D. tilosensis* and *D. catalonica* are aligned with mafft `(-einsi)`, an algorithm suitable to align conserved motifs (exons) embedded in an unalignable region (intron). Therefore, we are using clear homologous exons as “anchors” to improve the alignment of introns. *D. scabricula* sequences are added afterwards with mafft `(-add)` (Figure SM2g).

*D. scabricula* intron sequences were obtained using the same mapping approach described in section 1.1. Therefore, the consensus sequence obtained has the same coordinates as the reference but with the DNA information of *D. scabricula* (i.e. called variants in the mapped regions and Ns in the unmapped regions). Thus, if some region is missing in the reference sequence used when mapping, it will be also missing in the consensus sequence obtained (since there will be no region in the reference to map the read on). As a result, we are unable to distinguish between deletions in

*D. catalonica* and insertions in *D. tilosensis*. To address this issue, we generated two consensus sequences: i) mapping *D. scabricula* reads against *D. catalonica* (hereafter Dsca_1) and ii) mapping *D. scabricula* reads against *D. tilosensis* (hereafter Dsca_2). By leveraging the information from both consensus sequences (Dsca_1 and Dsca_2) we can discern between "apparent" insertions in *D. tilosensis* and deletions in *D. catalonica* (see Figure S2h for a schematic representation).

*1.5 Insertion/Deletions count*

First, we transformed pairs of aligned intronic regions to CIGAR (Concise Idiosyncratic Gapped Alignment Report) format (Table SM1).

- CIGAR 1: Indels in *D. tilosensis* compared to *D. catalonica*
- CIGAR 2: Indels in *D. tilosensis* compared to *D. scabricula* (Dsca_1)
- CIGAR 3: Indels in *D. tilosensis* compared to *D. scabricula* (both Dsca_1 and Dsca_2)

Second, we parsed CIGAR 1 string to identify stretches of deletions (D) in *D. tilosensis,* compared to *D. catalonica.* and also the same region for CIGAR 2, which allows us to consider outgroup information. We accept a region as a deletion when the studied region: i) has at least 70% 'D' characters, ii) is flanked upstream and downstream by 10 characters with at least 50% of mapped positions (M or = characters).

Similarly, we parsed CIGAR 1 to detect stretches of insertion (I) in *D. tilosensis*, and checked its region with CIGAR 2 information. In this case, we performed an additional check against CIGAR 3: if there is an insertion in *D. tilosensis*, then we should expect to observe a stretch of missing data (N, P, X) in the same region (at least 70%). In contrast, if there is a deletion in *D. catalonica*, we should expect to observe mapped positions (M or = characters). (Figure S2f). All these analyses were performed using custom Python scripts.

| **Reference (*D. tilosensis*)** | **Query** | **CIGAR character** |
|---|---|---|
| Nucleotide | Nucleotide | = for same nucleotides<br>M for different nucleotides |
| - | Nucleotide | D |
| Nucleotide | - | I |
| N (or -) | - (or N) | X |
| N (or nucleotide) | nucleotide (or N) | N |
| N | N | N |
| - | - | P |

Table SM1. CIGAR table of characters.

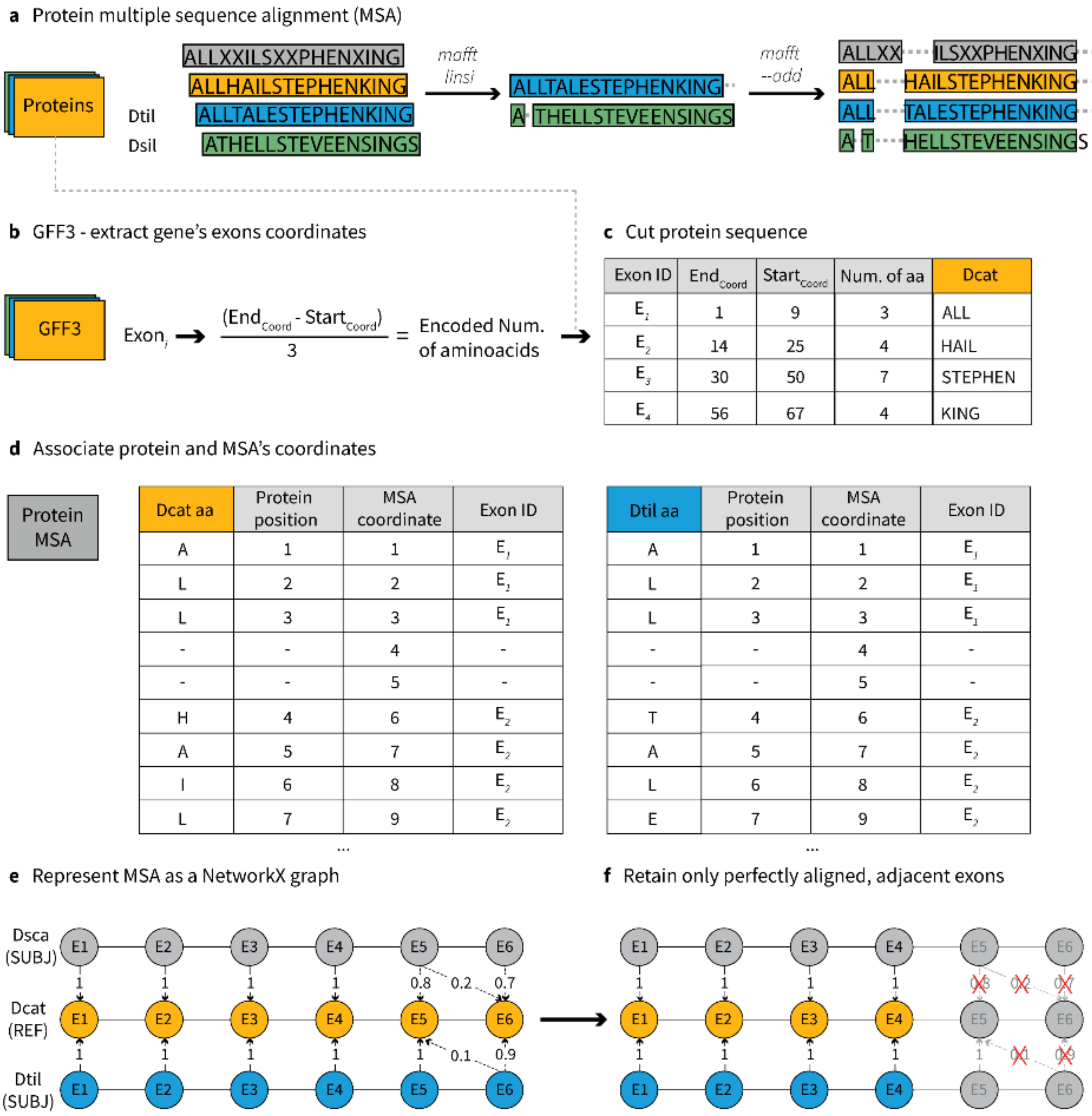


Figure SM2. Workflow for the indels identification in *D. tilosensis.*

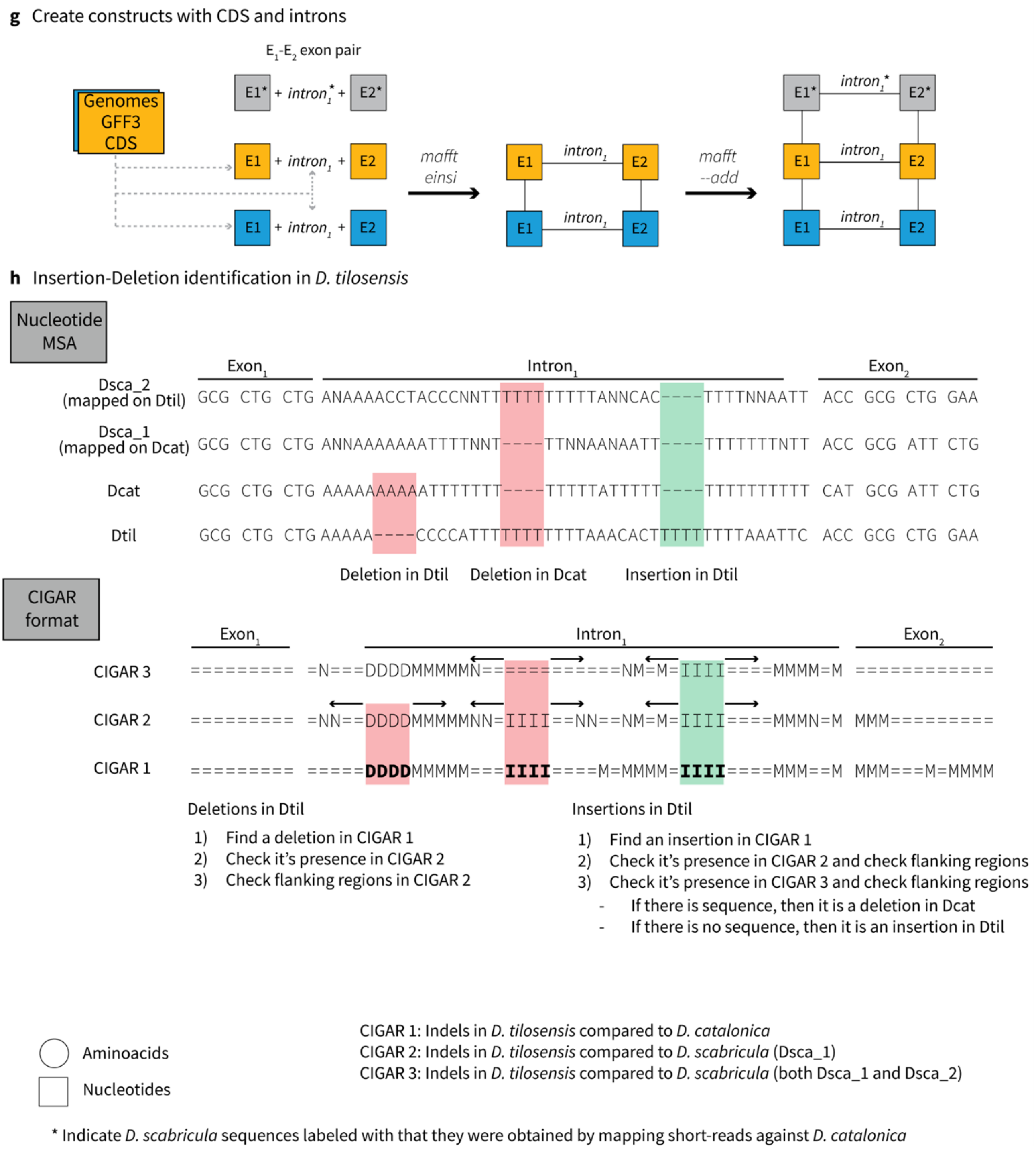


Figure SM2 (continued). Workflow for indels search in *D. tilosensis*.

### 2. Assessing the indels patterns in large intergenic data

We studied the indel patterns across large chromosomal fragments of *D. catalonica* and *D. tilosensis* using the mapped (aligned) information obtained with minimap2-2.24 (Li. H 2018). For the analysis, we only used information of the large scaffolds (chromosomes). We performed two mapping analyses: A1) *D. tilosensis* (query) against *D. catalonica* (target), and A2) the reciprocal, *D. catalonica* (query) against *D. tilosensis* (target). For this unpolarized analysis, we assume that the genome size at the ancestral *D. catalonica-tilosensis* split is around 3 Gb and, consequently, most indel events are presumed to be deletions occurring along the insular lineage leading to *D. tilosensis* (see also supplementary table 14).

Specifically, under A1, (a) Unmapped regions in the target genome (*D. catalonica*) are considered as deletions, and (b) Unmapped regions in the query genome (*D. tilosensis*) are considered as insertions. Under A2, (c) Unmapped regions in the target genome (*D. tilosensis*) are considered as insertions and, (d) Unmapped regions in the query genome (*D. catalonica*) are considered as deletions.

### 3. Recombination and nucleotide diversity analysis

For the three species (*D. catalonica*, *D. tilosensis* and *D. silvatica*), we estimated the per-site nucleotide diversity (π) and the population recombination rate (ρ) and examined the correlation between π and ρ, separately for autosomes and the X chromosome. We estimated π with VCFtools-v0.1.16 (Danecek et al. 2011) with option (`--site-pi`), and ρ values using the iSMC-v0.0.25 software/program (Barroso et al. 2019).

The raw reads from three resequencing experiments, one for each species (about 40x illumina PE 150) were trimmed in Trimmomatic v0.39 (Bolger et al. 2014) and mapped to their respective reference genomes using bwa mem v0.7.17-r1188 (Li and Durbin 2009) and short split hits labeled as secondary alignments (`-M`). Samtools v1.6 (Danecek et al. 2021; Li, et al. 2009) was used to retain only properly paired reads (`-f 2`), and filter out secondary alignments (`-F 256`) and alignments with MAPQ < 20 (`-q 20`). Picard Tools v2.26.10 (Broad Institute 2016) was used to remove duplicates, add read groups labels, index it and sort. We called SNPs with GATK HaplotypeCaller (GATK v4.3; Van der Auwera and O'Connor 2020) with option (`--min-base-quality-score 20`), and used GATK SelectVariants with options (`--select-type SNP, -select "DP > 10 && DP < 60"`) and GATK VariantFiltration to filter for QUAL < 30.0, QD < 2.0, SOR > 3.0, FS > 60.0 and MQ < 40.0.

We performed two complementary analyses, across the whole genome through a windows analysis (3.1), or in well-characterized genomic regions (i.e., syntenic regions, see below step 3.2). Statistical significance was assessed using the non-parametric Kendall rank correlation coefficient τ.

*3.1 Nucleotide polymorphism and recombination parameter in 1Mbp windows across genome*
The analysis was performed (separately for each of the three species and chromosomes) in 1 Mbp windows.

*3.2 Nucleotide polymorphism and recombination parameter in well characterized genomic data*
The analysis was performed only in syntenic regions between *D. catalonica* and *D. tilosensi*s. For that, we used information of the 1:1 orthogroups (7,547 cases), excluding information from small scaffolds, and considering only syntenic gene pairs (6,451 cases). We estimated the nucleotide diversity (π) and recombination rate (ρ) values only in these regions.
